# Investigating the impact of the interstitial fluid flow and hypoxia interface on cancer biology using a spheroid-on-chip perfusion system

**DOI:** 10.1101/2024.08.21.608919

**Authors:** Emily Pyne, Mark Reardon, Martin Christensen, Pablo Rodriguez Mateos, Scott Taylor, Alexander Iles, Ananya Choudhury, Nicole Pamme, Isabel M. Pires

## Abstract

Solid tumours are complex and heterogeneous systems, which exist in a dynamic biophysical microenvironment. Conventional cancer research methods have long relied on two-dimensional (2D) static cultures which neglect the dynamic, three-dimensional (3D) nature of the biophysical tumour microenvironment (TME), especially the role and impact of interstitial fluid flow (IFF). To address this, we undertook a transcriptome-wide analysis of the impact of IFF-like perfusion flow using a spheroid-on-chip microfluidic platform, which allows 3D cancer spheroids to be integrated into extracellular matrices (ECM)-like hydrogels and exposed to continuous perfusion, to mimic IFF in the TME. Importantly, we have performed these studies both in experimental (normoxia) and pathophysiological (hypoxia) oxygen conditions. Our data indicated that gene expression was altered by flow when compared to static conditions, and for the first time showed that these gene expression patterns differed in different oxygen tensions, reflecting a differential role of spheroid perfusion in IFF-like flow in tumour-relevant hypoxic conditions in the biophysical TME. We were also able to identify factors primarily linked with IFF-like conditions which are linked with prognostic value in cancer patients and therefore could correspond to a potential novel biomarker of IFF in cancer. This study therefore highlights the need to consider relevant oxygen conditions when studying the impact of flow in cancer biology, as well as demonstrating the potential of microfluidic models of flow to identify IFF-relevant tumour biomarkers.

## Introduction

The biophysiological forces present within the tumour microenvironment (TME) have gained increased interest for their roles in tumour progression in recent years. For example, there is extensive evidence that low oxygen in tumours (hypoxia), as well as nutrient provision and interactions with other cells, drives changes in gene expression, underpinning tumour progression and spread ^1-3^. Another aspect of the tumour biophysical TME is interstitial fluid flow (IFF), which has largely been overlooked in *in vitro* recapitulations of the TME. IFF velocity *in vivo* in tumours is generally accepted as being in the range of 1 × 10^−4^ – 10 mm s^-1^, although it is still poorly characterised ^4-6^. Of particular interest in the field is the impact it may have on the epithelial to mesenchymal transition (EMT) and early changes in tumour progression ^7, 8^. A large proportion of studies have focused specifically on the role of IFF in cell invasion and motility and many have been conducted in single cell models rather than 3D structures. One such study showed that IFF can guide tumour cell invasion along the flow direction ^9^. IFF has even been shown to promote tumour cell invasion via stromal-cell-mediated matrix remodelling within the TME, demonstrating a relationship between IFF and the TME ^10^. Other groups have used methods such as computational modelling to examine the spatial gradients in interstitial fluid that influence the direction of tumour cells as they invade ^6^.

More extensive work has been done regarding the effect of fluid shear stresses on gene expression changes, including on endothelial cells in vascular networks. Helle and colleagues examined transcriptomic remodelling of endothelial cells induced by flow replicating blood flow-induced shear stress ^11^. Further studies using renal epithelial cells also established transcriptomic changes caused by variations in fluid shear stress that led to upregulation of TGF-β, MAPK, and Wnt signalling pathways ^12, 13^. Yang & Xu and Follain and colleagues also used low shear stress to analyse gene expression profiles of endothelial cells and found significant differences in their transcriptomes as a result of flow ^14, 15^. Whilst these studies did not use cancer cells or indeed even a tumour-like microenvironment, they point to the significant influence that fluid flow can have on gene expression in these cells.

Many of the studies on IFF in the context of cancer have used 2D models or single cell suspensions in 3D environments, which do not simulate the structural depth and physiology of the TME. Spheroids have a 3D structure that results in heterogeneous acquisition of nutrients and oxygen diffusion throughout the spheroid, similar to that seen in tumours, making them a useful *in vitro* model ^16^. While spheroids better address the 3D nature of the TME they are generally still housed in static conditions and therefore do not incorporate fluid flow such as that of IFF. Microfluidic approaches - and specifically spheroid-on-chip models - are well suited to bridge this gap and incorporate fluid perfusion to model IFF, for example. However, while many studies have started to use lab-on-chip devices to study the TME, few have incorporated multiple, major critical components governing a tumour: IFF (continuous perfusion of media or fluid shear stresses), 3D structures (multicellular spheroids), and other relevant TME conditions such as hypoxia. We therefore set out to build on our lab’s spheroid-on-chip models ^17^ and expertise to perform an unbiased analysis of the impact of IFF on cancer biology in experimental (tissue culture) and, more importantly, tumour-relevant physiological (hypoxia) conditions, by conducting transcriptomic analyses to elucidate possible mechanisms by which IFF could be influencing TME progression. Our study is the first to identify that IFF-like flow-mediated gene expression changes in normoxic vs hypoxic conditions, as well as presenting a novel potential prognostic and predictive biomarker gene for IFF in cancer.

## Materials and Methods

### Microfluidic device design and fabrication

The device used in this study is a refined design of our previously published design ^17^. The updated design includes an inlet channel, a spheroid microwell with a removable polydimethylsiloxane (PDMS) plug seal, and an outlet channel (design details in **Figure 1A**). The microwell is 5 mm and 6 mm diameter for the lower chamber and top inlet hole respectively, which creates a step to prevent the PDMS from being pushed in too far into the well. Where channels interfaced with the central well, a teardrop shape was used for the junction (**Figure 1A, Supplementary Figure S1**). A 1 mm circular recess at the bottom of the well enabled a microscope cover slip to be positioned for spheroid imaging. Devices were fabricated by milling two glass slides (Schott B270) using a Computer Numerical Controlled (CNC) machine (M7 Datron, Germany) with a diamond milling tool of 1 mm diameter (Eternal Tools, UK). These were thermally bonded, and a borosilicate glass coverslip was subsequently bonded to the bottom of the device for the viewing port.

**Figure 1.**
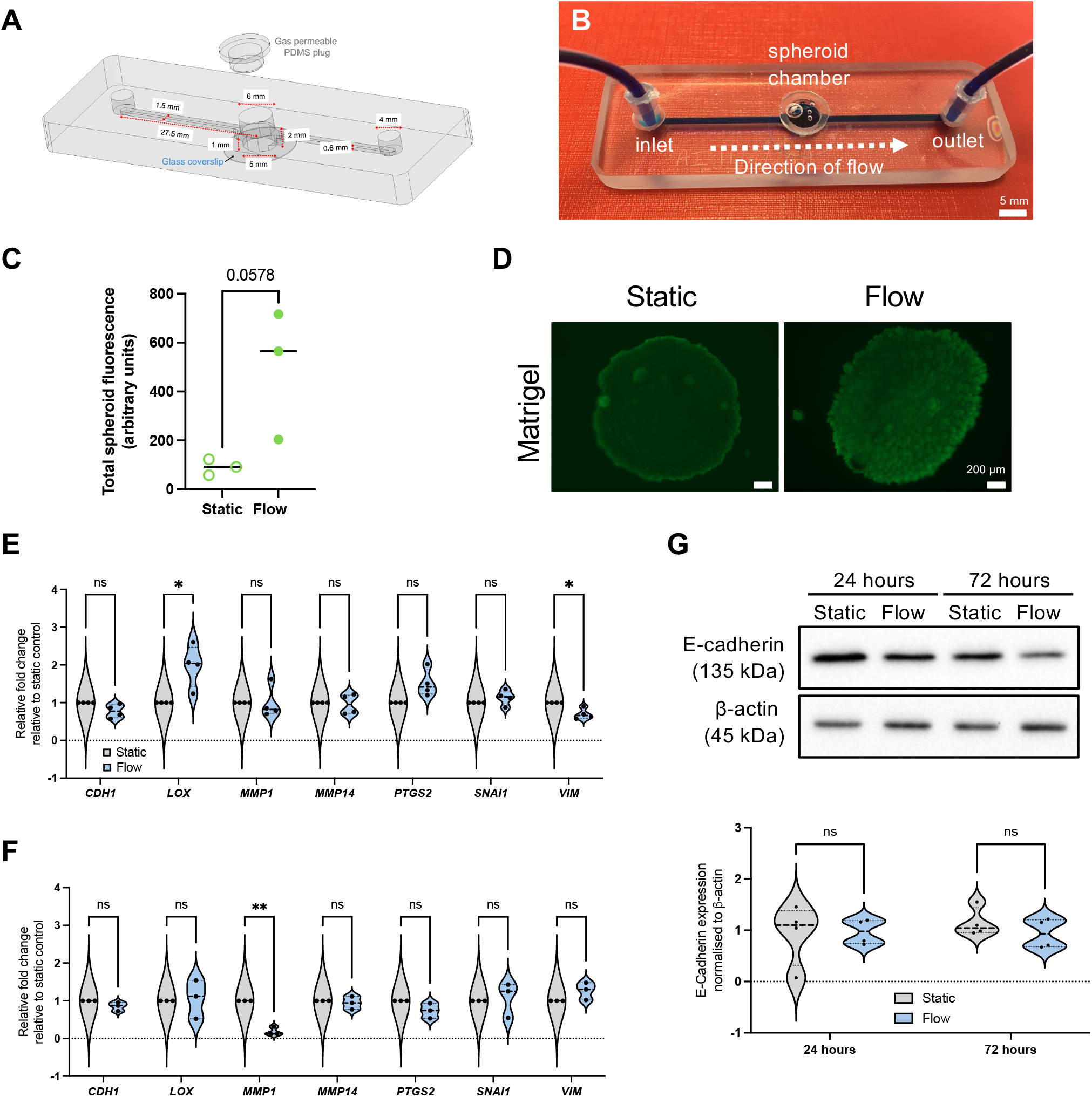
Updated spheroid on chip perfusion device design and initial evaluation of impact of IFF-like flow on EMT. (A-B) Schematic (A) and photograph (B) of the spheroid-on-chip microfluidic perfusion device showing the main components and key dimensions. (A) 4 mm wide inlet and outlet ports are placed at each end of the chip, with a 1.5 mm wide channel running from the inlet to the outlet. 27 mm downstream of the inlet, a spheroid chamber is included, with an access port with a PDMS plug for spheroid loading. (B) Brilliant blue dye flowing through the chip demonstrates the location of the channel and direction of flow. (C-D) Spheroids remain viable in flow conditions. MCF7 spheroids were formed from 9×10^4^ cells seeded in a ULA plate. Spheroids were then either kept on the ULA plate (Static) or transferred to the device (Flow; 3 μL min^−1^) for 72 hours in Matrigel. At the end of experiments, spheroids were stained with FDA and imaged. Dot plot represents average spheroid fluorescence (a.u.) and independent replicates. (D) Representative FDA staining images of MCF7 spheroids at 72 h incubated in static and flow conditions in Matrigel. Scale bar represents 200 μm. To determine impact of IFF-like flow, MCF7 spheroids were formed from 9×10^4^ cells as before, and exposed to in static or flow conditions for 24 or 72 hours in the presence of Matrigel. Total RNA (E-F) or protein samples (G) were extracted and qPCR (E-F) or immunoblotting (G) performed. For qPCR (E-F), a panel of EMT relevant genes was analysed, with *B2M* used as housekeeping gene. Violin plots represent n=4 (E; 24 hours) or n=3 (F; 72 hours) independent experiments. Immunoblotting was used to analyse E-cadherin protein expression levels (G). Densitometry was used to quantify bands, and band intensity relative to loading control depicted in the violin plot. Blots are representative of n = 3 experiments. Unpaired student’s t-test was performed to test for statistical significance between samples. ns = nonsignificant; * p<0.05; ** p<0.01

### Microfluidic spheroid-on-chip setup

Spheroids were generated from the MCF7 (pleural effusion of metastatic breast adenocarcinoma) cell line by seeding 9×10^4^ cells per well in ultra-low adherence (ULA) plates. Cells were purchased from authenticated source European Collection of Authenticated Cell Cultures (ECACC), and regularly tested negative for mycoplasma.

A spheroid was transferred to the chip microwell in 50 µL of media. Chilled pipette tips were used to add 50 µL of Matrigel and it was allowed to set before the chamber was sealed with a PDMS plug. The device was attached to a syringe pump (Harvard Apparatus). A plastic syringe (BD, Oxford, UK) filled with complete Dulbecco’s Modified Eagle Medium (DMEM) was secured in the syringe pump and attached to Tygon tubing (Cole-Parmer, Tygon Microbore), 1.6 mm inner diameter with a 200 µL pipette tip, which had been cut to fit securely over the syringe nozzle (**Figure 1B**). At the outlet end of the chip, Tygon tubing was used as before, leading to a collection pot for effluent media. Flow rate was set at 3 μL min^−1^, and velocities were 1.49 mm min^-1^ (tubing), 3.3 mm min^-1^ (channels), and 0.2-0.6 mm min^-1^ (microwell, as specified later).

### Hypoxic experiments

A H35 Hypoxistation hypoxia chamber (Don Whitley Scientific, UK) was used for achieving hypoxic conditions of 1% O_2_. All equipment, media, and consumables were equilibrated overnight prior to experiments.

### FDA viability assay

Fluorescence live staining assays using fluorescein diacetate (FDA) (Thermo Fisher Scientific) were conducted at the end of the experiment as previously described ^17^. The fluorescent signal produced by FDA was quantified using ImageJ ^18^ with a method adapted from McCloy er al ^19^. The ‘area’, ‘integrated density’ and ‘mean grey value’ options were selected in the ImageJ ‘analyse’ tool bar menu. Using greyscale images, the ‘polygon’ tool was then used to draw around the spheroid as the region of interest (ROI). A background signal measurement located away from the ROI was taken at the same size as the ROI. To produce the corrected total spheroid fluorescence (CTSF), the background fluorescent signal was subtracted from the integrated density value, and this was represented as relative levels between two conditions.

### Quantitative real-time PCR (qRT-PCR)

RNA extraction and qRT-PCR was performed as previously described ^20^. In brief, RNA was extracted using the Aurum Total RNA mini Kit (Biorad, UK) and relative mRNA expression from static and flow spheroids using QuantiFAST SYBR Green (Qiagen) and the StepOnePlus™ Real-Time PCR System (Thermo Scientific). Transcript expression levels were normalised to *B2M* (β-2-microglobulin). Primer details are available in **Supplementary Table S1**. Mean CT values of independent biological replicates (see Figure legends for specific replicate numbers) were used to calculate relative expression of gene targets using the 2^-ΔΔCT^ method ^21^.

### Immunoblotting

Spheroid lysates were prepared in UTB (Urea-Tris-Beta-mercaptoethanol) buffer and analysed by immunoblotting as previously described ^22^. Antibody details are available in **Supplementary Table S2**. Band intensity densitometric quantification was performed using Image J (NIH) ^18^.

### RNA-sequencing and bioinformatic analysis

Spheroids were collected from three biological replicates of either static (ULA plate) or flow (perfusion chip) conditions for RNA extraction as previously described. mRNA library preparation (using poly A enrichment) and sequencing was performed by Novogene (Cambridge, UK), using an Illumina NovaSeq PE150 (paired end, 150 bp read length) platform with ≥ 20 million read pairs per sample, using HISAT2 for mapping and FPKM (Fragments Per Kilobase of transcript sequence per Millions base pairs sequenced) for gene expression estimation. Differential expression analysis was performed from read counts between the two conditions (static and flow) with three biological replicates for each, using the DESeq2 R package ^23^. FPKM-derived, differentially expressed gene expression levels were transformed by adding one (to avoid taking the logarithm of zero), taking the binary logarithm (log_2_) and subtracting the genes’ medians. DESeq2-derived p-values were corrected for multiple testing with the Benjamini and Hochberg method ^24^. Differentially expressed genes (DEG) were filtered to those with adjusted p-value < 0.05 and absolute fold change > log_2_(1.5). DEG heatmaps were generated with R package ComplexHeatmap v2.16.0 ^25^. DEG volcano plots were generated with R package EnhancedVolcano v1.18.0 ^26^. R packages clusterProfiler v4.8.2 ^27^ and org.Hs.eg.db v3.17.0 ^28^ were used to perform Gene Ontology enrichment analyses and generate dotplots. All analyses were performed with R v4.3.0 ^29^. Canonical pathway diagrams were generated using Ingenuity pathway analysis (IPA; Qiagen). Datasets will be available on Gene Expression Omnibus (GEO) ^30^ once study is published.

### VEGF ELISA

Quantification of VEGF (Vascular Endothelial Growth Factor) in conditioned media and effluent was detected using a VEGF ELISA kit (Quantikine, R&D Systems) as per manufacturer’s instructions. Secreted factor levels were normalised as picograms (pg) per mL of media.

### COMSOL simulations

Computer simulations were carried out using COMSOL Multiphysics 5.2 (COMSOL Inc.) as before ^17^, with modifications. COMSOL was used to simulate the oxygen consumption in spheroids in the microfluidic system and in static conditions. A flow rate of 3 μL min^−1^ (as applied in all flow experiments) and a hypoxic threshold oxygen concentration of 10 mmHg (equivalent to 1.3% O_2_) were used. Simulations were performed using a model where reaction rate based on Michaelis–Menten model is assumed inside the spheroids, as noted in previous studies ^31, 32^.

### *In silico* patient dataset analyses

The prognostic value of candidate biomarkers was determined using the RNA-seq dataset for breast invasive carcinoma samples (n=1,110) ^33^ downloaded from the publicly available TCGA (The Cancer Genome Atlas) project accessed through cBioportal ^34-36^. cBioPortal was also used to evaluate the impact of ACTL8 expression for relevant clinical attributes, including breast cancer subtypes and hypoxia scores ^37, 38^. The online tool KMplot ^39^ was used to determine the prognostic value of specific genes for an additional breast cancer dataset (Metabric dataset) as well as other tumour types in the pan-cancer TCGA datasets (see **Supplementary Figures S6** and **S9** respectively for more details). For these analyses, median gene expression was used to split patient cohorts into groups with “high” and “low” expression. The online tool ROCplotter was used for evaluation of a gene of interest as a predictive biomarker for breast cancer ^40^. The tool generated an ROC (receiver operating characteristic) plot with AUC (area under the curve) computed to assess prognostic power.

### Statistical analysis

All experiments were performed independently at least three times (biological replicates). Paired t-tests were used to compare two groups (*e*.*g*. static and flow conditions), and the Holm-Šídák method was used for correction when multiple t-testing was required. Where more than two groups were compared with one another, a two-way ANOVA followed and Kruskal-Wallis testing was used. For Kaplan-Meier survival plots, hazard ratios with 95% confidence intervals and logrank P values were calculated. ROC significance was performed using the Mann-Whitney test. Statistical analyses were performed using GraphPad Prism 10.2.3 (California, USA).

## Results and Discussion

### Updated perfusion device design and initial evaluation of IFF impact on spheroid on chip

Previous work in our lab developed a microfluidic device capable of housing viable spheroids for up to 72 h in continuous perfusion to study the effects of flow ^17^. We have further refined this design to improve spheroid housing, including a larger microwell, as well as a more streamlined design (**Figure 1A-B**), including even volume channels and a teardrop shape used for the junction between microwell and outlet (**Supplementary Figure S1**). This redesign led to the ability to establish velocities and shear stresses more relevant to reported IFF-relevant speed parameters ^7^. Flow rates utilised for these experiments were setup at 3 µL min^-1^, which led to a median residence time in the spheroid microwell of 27 minutes, and shear stress of 5.64 × 10^−5^ (Matrigel) or 6.27 × 10^−6^ (media alone) Dyne cm^-2^, velocity of 0.6 (Matrigel) and 0.2 (media alone) mm min^-1^, and a Reynolds number of 1.79 × 10^−2^ (Matrigel) or 1.34 × 10^−2^ (media alone), indicating laminar flow.

As in our previous study ^17^, we determined spheroid viability using FDA *in situ* viability assay, and it can be observed that, albeit not significantly, spheroid viability is increased under flow when compared with static conditions, as shown by an increase of FDA associated fluorescence (**Figure 1C-D**).

As noted earlier, there are several studies using 2D or 3D single cell models showing that high flow or perfusion rates leads to increased expression of ECM remodelling and EMT markers ^8, 41, 42^. T herefore, we used the epithelial-like cell line MCF7 to perform did an initial evaluation of the expression of some key EMT (E-Cadherin - *CDH1*; Vimentin - *VIM*; Snail - SNAI1; COX2 - *PTGS2*) and ECM remodelling (Lysyl Oxidase - *LOX*, Metalloproteinases *MMP1* and *MMP14*) markers (**Figure 1 E-G**). However, surprisingly, most of these factors were not significantly altered, and some were even downregulated. It would be interesting to use even less tumour-like models, such as spheroids derived from non-tumour immortalised cell lines such as PrEC-hTERT (prostate) and MCF10A and HMEC (breast). Only in one other study that used 3D “tumouroids” (cells aggregated into compact masses) was the same lack of upregulation of EMT markers observed. In this study, tumouroids were exposed to both fluid flow and pressure as mechano-inducing stresses in a microfluidic system ^43^. Importantly, they did not see overall increases in EMT markers either and in fact reported decreased expression of vimentin. Notably, they did observe a significant increase in MMP14 transcript, which contrasts with our results. Also, as well as the induction of pressure, the authors included collagen rather than Matrigel and used a more mesenchymal cell line (MDA-MB-231) rather than the more epithelial-like model (MCF7) used in our study. It would be interesting to compare the impact of flow alone to the combination of flow and pressure on gene expression using comparable experimental conditions.

These data indicated that, at least for our model, flow/perfusion rates associated with IFF did not induce upregulation of markers of EMT and early invasion, contrary to most previously reported studies. Therefore, we hypothesised that flow/perfusion might regulate other cancer-related pathways at IFF-relevant flow rates.Whole transcriptome analysis reveals impact of IFF-like perfusion on cancer spheroid biology, including upregulation of cell cycle and DNA repair pathways

In order to perform an unbiased analysis of gene expression changes associated with perfusion or flow at IFF-relevant rates, we performed whole-transcriptome RNA-sequencing (RNA-Seq) analysis, to provide us with a broader snapshot of differentially expressed genes (DEGs) under flow. For this, total RNA samples from MCF7 spheroids exposed to static or flow conditions for 24h were analysed using RNA-seq (**Figure 2**). The comparison between the transcriptomes of static and flow MCF7 spheroids showed significant differences in overall expression patterns evident between the two conditions, as can be observed in the relevant DEGs heatmap (**Figure 2A**) and volcano plot (**Figure 2B**).

**Figure 2.**
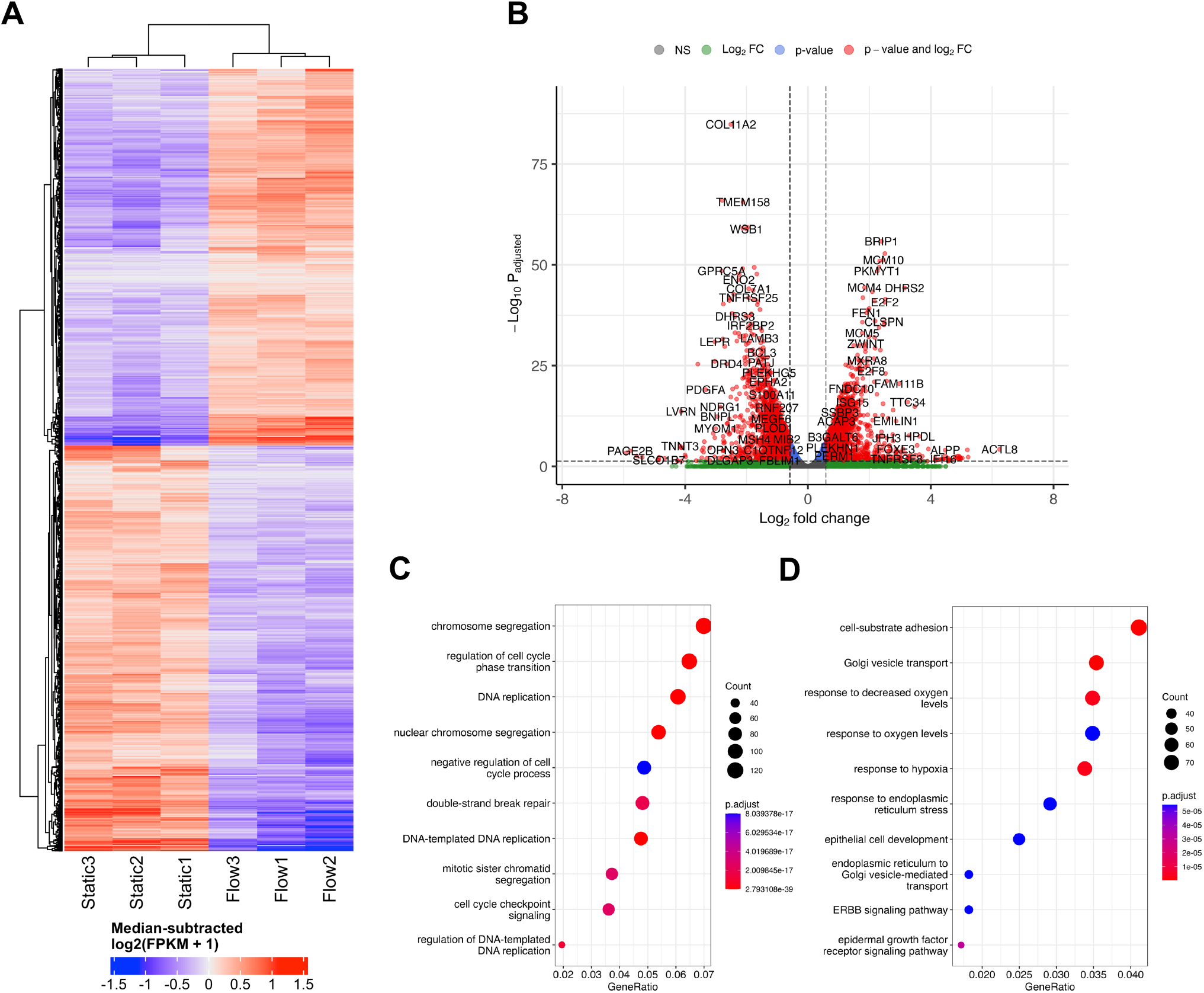
Whole transcriptome analysis reveals impact of IFF-like perfusion on cancer spheroid biology. To determine the whole transcriptome impact of IFF-like flow, MCF7 spheroids were formed from 9×10^4^ cells as before, and exposed to static or flow conditions for 24 h in the presence of Matrigel. Total RNA was extracted, and bulk RNA-sequencing was performed on three independent samples per condition. (A-B) Heatmap (A) and volcano plot (B) represent significantly differentially expressed genes (DEGs), with p.adjust<0.05 and fold change > log_2_(1.5). Green: genes whose expression are above the Log_2_ FC threshold; blue: genes whose expression are above the p-value threshold; Red: genes whose expression are above both p-value and Log_2_ FC thresholds (red). (C-D) Pathway enrichment analysis using GO (gene ontology) categories was performed for upregulated (C) and downregulated (D) genes in flow when compared with static conditions.

Gene Ontology (GO) canonical pathway enrichment analysis was performed for both upregulated (**Figure 2C**) and downregulated (**Figure 2D**) DEGs in flow vs static conditions. The top canonical pathways and functions upregulated in the differential expression between static and flow were heavily represented in the categories of cell cycle/DNA replication and DNA repair processes. Conversely, the top canonical pathways in the downregulated cohort included endoplasmic reticulum (ER) stress and Golgi transport, as well as adhesion, and response to hypoxia/oxygen levels. It is plausible that pathway enrichment might be dependent on the cell line model used and flow rates used, and further work needs to be performed elucidate this. For example, a recently published study looked at culturing primary patient-derived glioblastoma (brain cancer) cell lines in static *vs* flow conditions (shear stress 4.9 × 10^−5^ Dyne cm^-3^) and evaluated whole transcriptome gene expression changes ^44^. They showed enrichment for cell migration, ECM interaction, metabolism and cell death, amongst others, which differs from our observations. It is important to note that this model defined static as pre-flow conditions, rather than a pairwise comparison with time in flow, as we use in our system. Similarly, Goetz’s group has previously cultured human primary endothelial cells (HUVEC) in static and flow conditions in a microfluidic device to analyse the impact of flow on endothelial biology ^14^. Here, a whole transcriptome analysis revealed that these high flow rates relevant for vascular flow (400 µm s^-1^) led to dramatic changes in gene expression, but, unlike our study, cell cycle relevant factors were downregulated, with factors related to angiogenesis being upregulated. A more detailed comparison of the nuances between our, these, and other relevant studies, as well as an independent analysis using comparable models, is necessary to have a comprehensive understanding of the nuances of the impact of different physiologically relevant flow conditions in various models.

### IFF-like perfusion/flow led to upregulation of cell cycle and DNA repair pathways

To validate our observations, we initially focused on pathways identified as enriched for upregulated DEGs to be validated, namely relevant to DNA damage response (DDR) pathways: DNA replication/cell cycle and DNA repair (**Figure 3, Supplementary Figure S2**). For DNA replication and cell cycle regulation we evaluated the expression of *MCM6* (Minichromosome Maintenance Complex Component 6), *RRM1* and *2* (Ribonucleotide Reductase Regulatory Subunits M1 and 2), *ASF1B* (Anti-Silencing Function 1B Histone Chaperone), *WEE1*, and *E2F1*, (**Figure 3 A-B**) and for DNA repair we evaluated expression of *BLM, BRCA1, BRCA2, EXO1* (Exonuclease 1), *FEN1* (Flap Structure-Specific Endonuclease 1), and *RAD51* (**Figure 3 C-D**). These data show that, overall, most of the selected factors were validated independently, and that the increased expression of these factors was sustained up to 72 h, indicating the observed changes are not solely due to an acute response to better perfusion. However, to fully evaluate the impact of perfusion with well-oxygenated media of a spheroid mass which was setup to include oxygen gradients and therefore is partially hypoxic, analysis of markers for DNA damage and replication stress needs to be completed.

**Figure 3.**
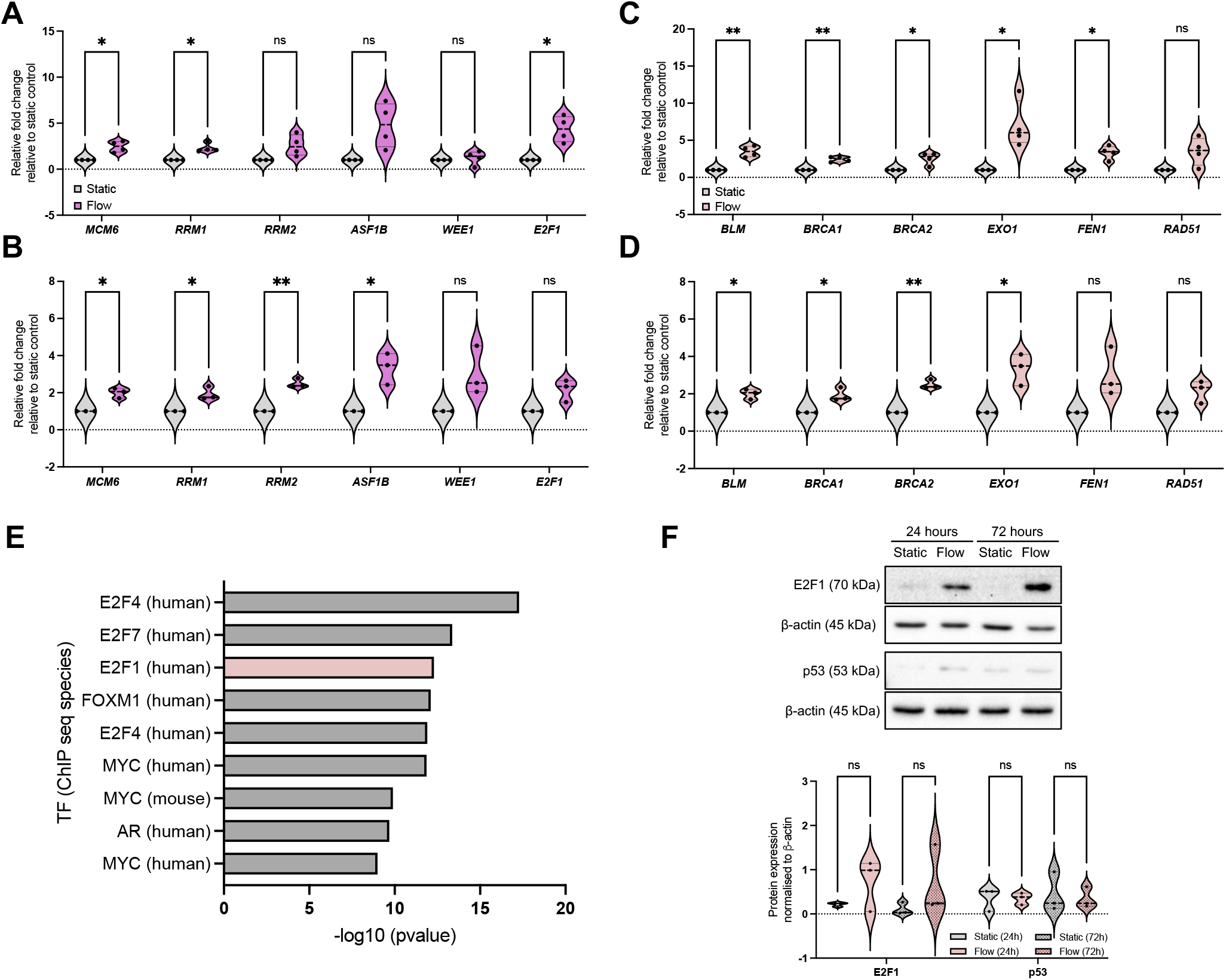
IFF-like perfusion/flow led to upregulation of DNA replication, cell cycle, and DNA repair pathways. (A-D) DNA replication/cell cycle (A-B) and DNA repair (C-D) signalling pathways were enriched in upregulated DEGs after flow. A panel of genes for these pathways was validated through qPCR as described before, with plots representing n=4 (A, C; 24 h) or n=3 (B, D; 72 h) independent experiments. (E) Transcription factor (TF) enrichment was analysed for the upregulated DEGs. Immunoblotting (F) was used to analyse the protein expression of E2F1 and p53. Densitometry was used to quantify bands, and band intensity determined relative to loading control. Blots are representative of n = 3 experiments. Unpaired student’s t-test was performed to test for statistical significance between samples. ns = nonsignificant; * p<0.05; ** p<0.01

To elucidate the potential mechanism underpinning these changes, we performed a transcription factor (TF) enrichment analysis on the upregulated DEGs (**Figure 3E**) and identified that TFs likely involved in these gene expression changes include the E2F family ^45, 46^. We further evaluated E2F1 protein expression under static and flow conditions at these timepoints (**Figure 3F**). Although we found that E2F1 protein expression increased as a trend, this was not statistically significant. Analysis of the E2F family might provide a clearer picture for their role in this process, as well as analysis of their activity using reporter assays ^47^. We also investigated the levels of p53, another key TF relevant for DDR-regulated gene expression regulation, and these were not significantly altered (**Figure 3F**).

### Hypoxia signalling is downregulated in the better-perfused spheroid-on-chip model

We next evaluated some of the top downregulated pathways from the enrichment analysis, focusing on response to hypoxia (**Figure 4, Supplementary Figure S3**). For this, we evaluated the expression of well-established target genes of the HIF (hypoxia-inducible factor) TF family, namely *CA9* (Carbonic Anhydrase 9), *BNIP3* (BCL2 Interacting Protein 3), *WSB1* (WD Repeat And SOCS Box Containing 1), and *VEGFA* (**Figure 4A and Supplementary Figure S3B**).

**Figure 4.**
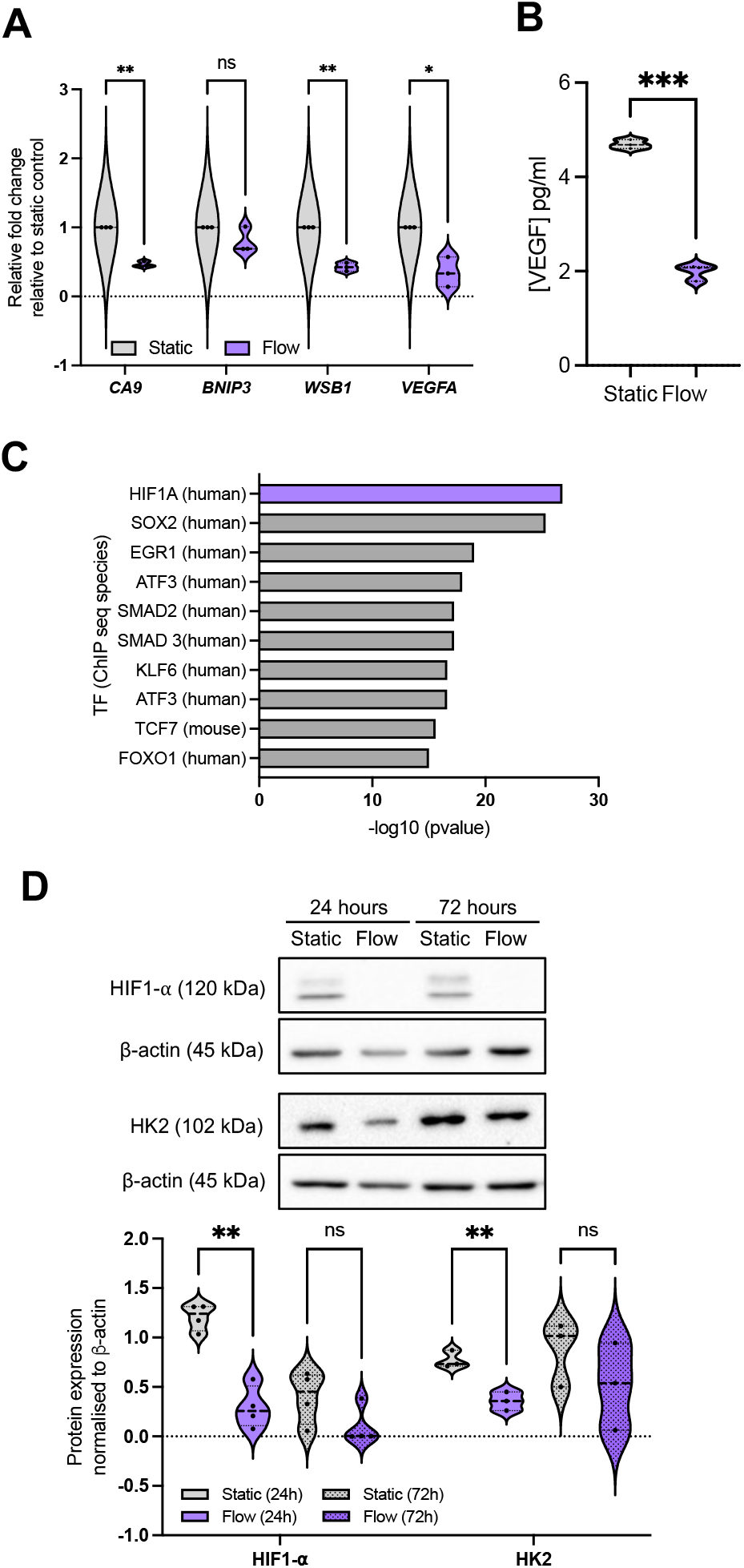
Hypoxia signalling is downregulated in the better perfused spheroid-on-chip model. (A) A panel of hypoxia-regulated genes was validated through qPCR as described before for 24 hours post flow, representing n=3 independent experiments. (B) Conditioned media and effluent were collected from these samples at 24 hours and VEGF concentration (pg mL^-1^ media) analysed by ELISA. (C) Data for 72 h samples is included in Supplementary Figure S3. Transcription factor (TF) enrichment was analysed for the downregulated DEGs. (D) Immunoblotting was used to analyse HIF1-α and HK2 (hexokinase 2) protein levels. Densitometry was used to quantify bands, and band intensity determined relative to loading control. Blots are representative of n=4 (24 h) or n=3 (72 h) experiments. Unpaired student’s t-test was performed to test for statistical significance between samples. ns = nonsignificant; * p<0.05; ** p<0.01; *** p<0.01

As expected, by improving spheroid perfusion with well-oxygenated media, we observed a decrease in HIF-related gene expression patterns after 24 hours in flow (**Figure 4A**), with the impact being reduced after 72 h in flow (**Supplementary Figure S3B**). This is reflected in the decrease in HIF1-α protein stabilisation under flow (**Figure 4D**). A similar pattern was observed for VEGF protein levels in the effluent media (**Figure 4B**). We again performed a TF enrichment analysis on the upregulated DEGs (**Figure 4C**) which confirmed HIF1-α as the top TF for the significantly downregulated DEGs. Other TFs were also highlighted (**Figure 4C**), which will be evaluated in further studies. This reduction in hypoxia signalling raised the question of whether the gene expression changes observed between static and flow conditions could be predominantly due to better spheroid perfusion and oxygenation, especially considering that chronic hypoxia can lead to repression of DNA repair, replication, and cell cycle regulatory factors ^47-49^. This is key to clarify for our model, as hypoxia is prominent in most solid tumours and it is closely linked with alterations in perfusion, IFF, and interstitial fluid pressure (IFP) due to the irregular and non-productive tumour vasculature ^50, 51^. Therefore, we decided to further investigate the interplay between these two TME components.

### Gene expression changes mediated by IFF-like perfusion/flow differ depending on environmental oxygen tensions

To elucidate this, we first modelled the impact on spheroid oxygen consumption if the media was differentially oxygenated using COMSOL, including atmospheric (20%), physoxic or physioxic (3%) and hypoxic (1%) oxygen levels (**Figure 5**), using a similar approach as in our previous study ^17^.

**Figure 5.**
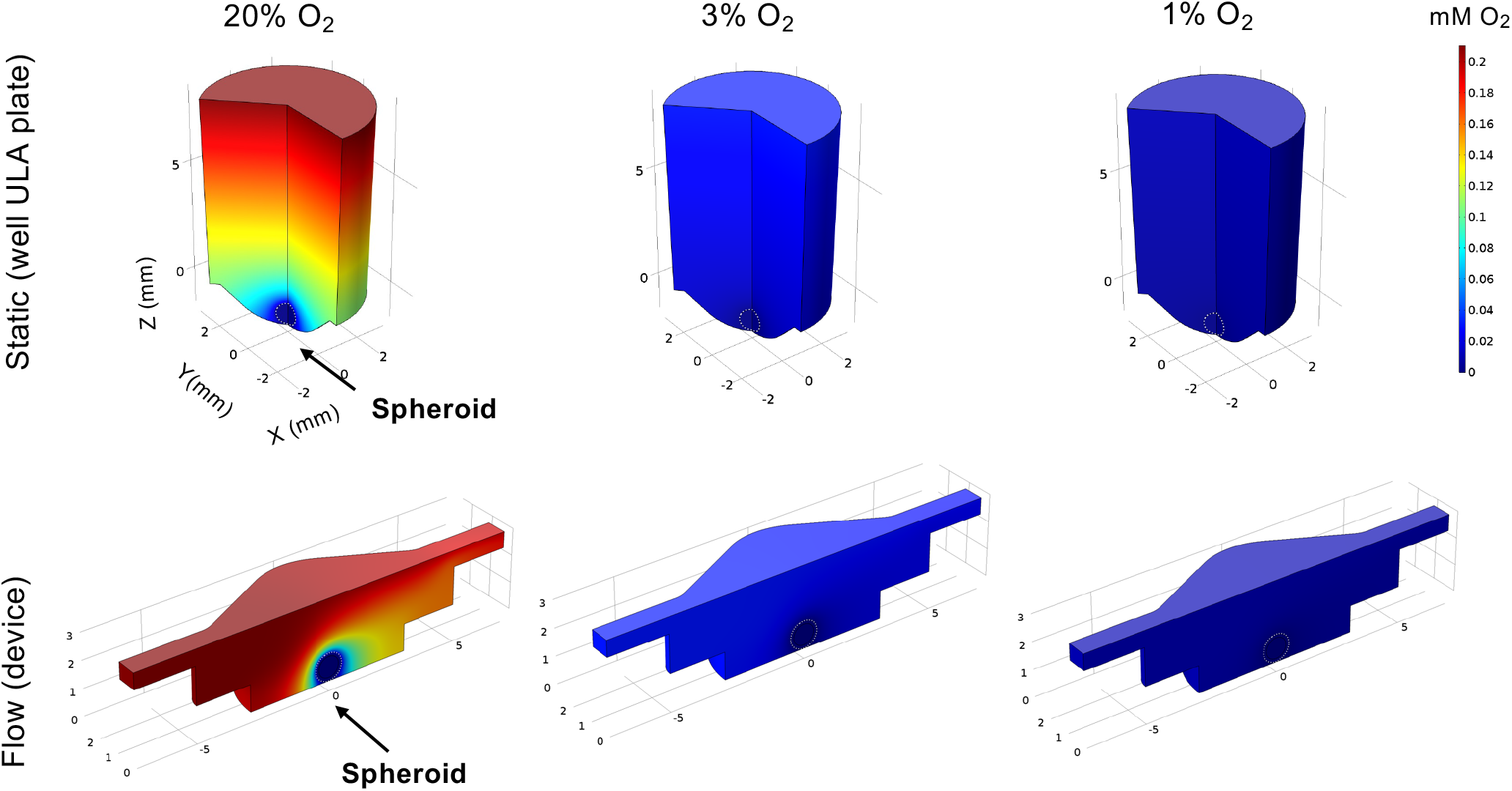
Spheroid cell oxygen consumption modelling for static vs IFF-like flow conditions in varying oxygen conditions. Oxygen consumption modelling of spheroids in static and flow conditions was performed, with the spheroid perfused with air saturated medium (no matrigel) at three different oxygen tensions (20%, 3%, and 1% O_2_ levels) in static (ULA plate) conditions or at a flow rate of 3 μL min^−1^. The mid-height cross-sectional line of the spheroid is indicated by the white dashed line. Oxygen levels colour gradients represent oxygen levels in mM O_2_.

We defined a hypoxic threshold as 1.3% O_2_ ^16^, using the predicted oxygen consumption rates as defined elsewhere ^31, 32^. As expected, cells in the spheroids are below the hypoxic threshold when media has 1% O_2_. However, if the media is better oxygenated, higher fractions of the spheroid mass are above the hypoxia threshold (0.4 for 20% O_2_ and 0.05 for 3% O_2_, respectively) in flow conditions, when compared to static conditions (0.2 for 20% O_2_ and 0 for 3% O_2_, respectively). This indicates that, to best represent the biophysical TME), both static and flow conditions should be fully at or below the hypoxic threshold of 1% O_2_.

We next conducted another whole-transcriptome RNA-seq analysis in low oxygen (1% O_2_) to evaluate DEG patterns in flow under more relevant hypoxic conditions. This was performed by placing both static plate and flow device setup in a specialised hypoxia chamber and collecting and analysing RNA samples for each condition as before (**Figure 6, Supplementary Figure S4**).

**Figure 6.**
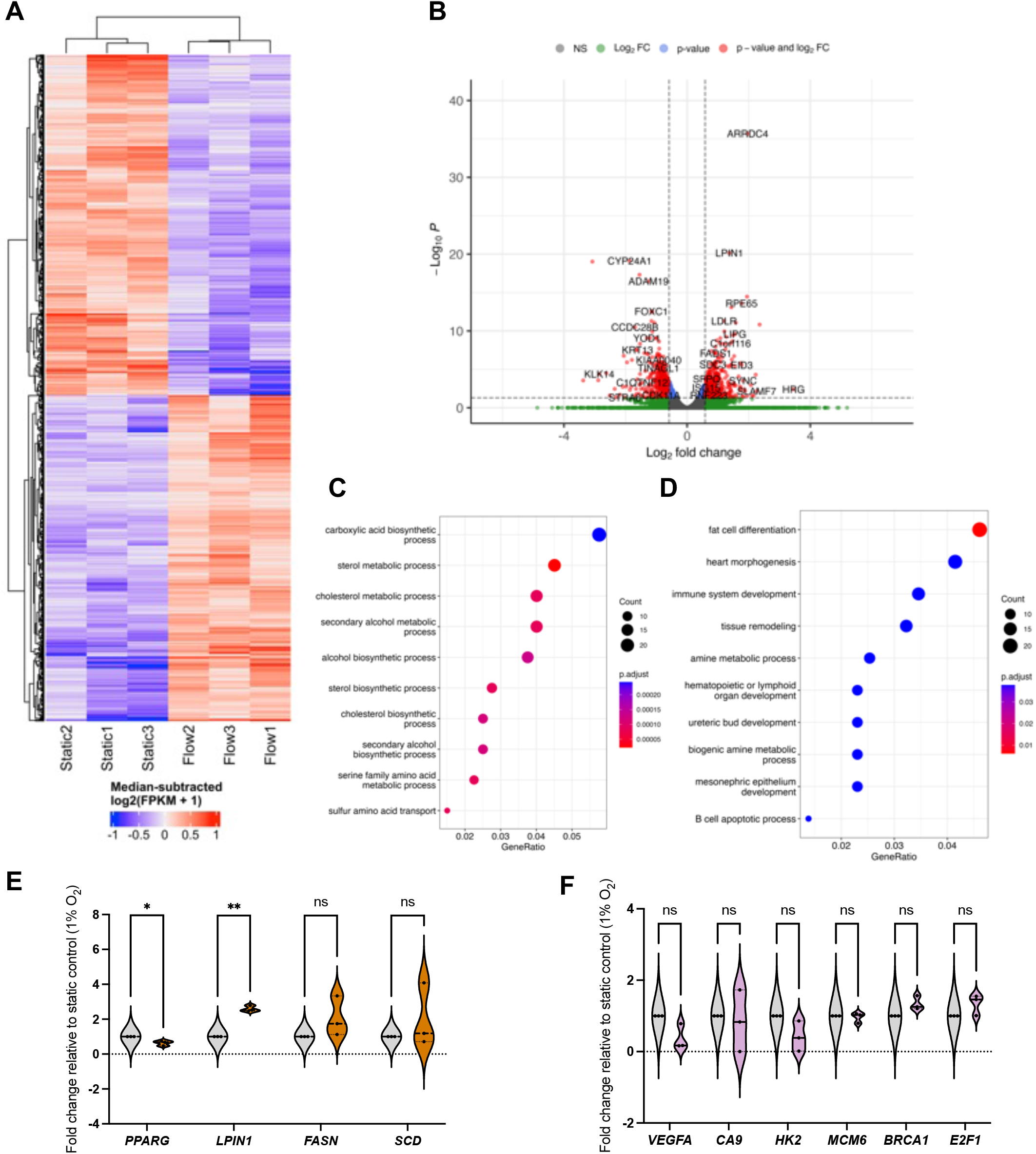
Gene expression changes mediated by IFF-like perfusion/flow differ depending on environmental oxygen tensions. To determine the whole transcriptome impact of IFF-like flow in hypoxic conditions, MCF7 spheroids were formed from 9×10^4^ cells as before, and exposed to in static or flow conditions for 24 h in the presence of Matrigel in a hypoxia chamber set at 1% O_2_. Total RNA was extracted, and bulk RNA-sequencing was performed on three independent samples per condition. (A-B) Heatmap (A) and volcano plot (B) represent significantly differentially expressed genes (DEGs), with p.adjust<0.05 and fold change > log2(1.5). (C-D) Pathway enrichment analysis using GO (gene ontology) categories was performed for upregulated (C) and downregulated (D) genes as before. (E-F) Phospholipid metabolism signalling pathways were enriched in upregulated DEGS after flow. A panel of genes for these pathways (E) and hypoxia and DDR (F) was validated through qPCR as described before, with plots representing n=3 (24 h) independent experiments. Unpaired student’s t-test was performed to test for statistical significance between samples. ns = nonsignificant; * p<0.05; ** p<0.01

This is, to our best knowledge, the first time transcriptome-wide analyses of the impact of IFF have been performed in low oxygen conditions relevant to the biophysical TME. As observed in normoxia, there are clear differences between static and flow samples in terms of gene expression patterns (**Figure 6A-B**). Interestingly, the GO canonical pathway analysis revealed different patterns from those observed in normoxia (**Figure 6C-F**). Specifically, metabolic pathways, including lipid and amino-acid metabolism, are enriched for upregulated DEGs, with development-linked pathways enriched in downregulated DEGs. Pathways altered by flow in normoxic conditions, such as DDR and hypoxia, are not significantly altered by flow in hypoxic conditions (**Figure 6F**).

Fatty acid metabolism is well established to be associated with tumour progression adaptation ^52^. Hypoxic conditions are associated with poor vascularisation and insufficient nutrient supply, including fatty acids (FAs). Flow conditions allow for a better perfusion of the spheroids with more nutrient-rich media than static conditions and could thus lead to increased uptake of FAs in flow *versus* static settings even in hypoxic conditions, potentially leading to the increased FA metabolism and other fatty acid metabolic pathways, and supporting *de novo* lipid synthesis ^52^. The FA-synthesis-associated enzyme *FASN* (fatty acid synthase) is upregulated, albeit not significantly, whereas LIPIN (*LPIN1*), linked with triglycerides and lipid droplets accumulation, was significantly upregulated in flow conditions (**Figure 6E**). It would be interesting to evaluate the impact of flow as a mechanical stimulus alone by using nutrient-depleted media in hypoxic conditions, and to evaluate whether these pathways remain upregulated. The exact role of lipid biology in interstitial-flow-regulated biology, and how the expression of these is regulated, will be evaluated in future studies. Interestingly we have observed the TF PPARγ (*PPARG*) to be downregulated in flow conditions (**Figure 6E**). PPARγ is a key regulator of metabolic remodelling, with reported tumour-suppressive functions in cancer, although somewhat controversially ^53^. Downregulation of PPARγ in flow further reiterates that, albeit in hypoxic conditions, flow-mediated perfusion of the spheroids with more nutrient-rich media would lead to a decrease in the need to rely on PPARγ-regulated pathways and, potentially, its downregulation compared to nutrient-deprived static conditions.

### Identification of potential IFF-associated clinically relevant biomarkers – a proof-of-principle analysis

Attempts have been made at using interstitial fluid as a sampling medium for tumour-specific proteins and other biomarker candidates, also referred to as the cancer secretome, for studying proteins secreted by cancer cells and other cells in the confined environment of a tumour ^54, 55^. However, there has been less work on establishing biomarkers linked with IFF. Interstitial fluid pressure (IFP) is in some cases considered a predictive biomarker, with an increase in IFP being linked with poor prognosis ^56-58^. However, analysis of molecular markers of IFP is again limited; to our knowledge no work has been done to specifically establish biomarkers of fluid flow. We hypothesized that identification of a gene or genes significantly differentially expressed in transcriptome-wide analyses could be assessed against various clinical outcomes to identify a potential biomarker of IFF in the TME. To do this, we identified three potential markers uniquely and significantly (p<0.01) expressed in either static or flow conditions, and with some indications in the literature of relevance to cancer biology: *ACTL8* (Actin Like 8), *RAB6C* (RAB6C, Member RAS Oncogene Family), and *BARHL1* (BarH Like Homeobox 1) (**Figure 7A**). Of these, only ACTL8 was independently validated in both 20% and 1% O_2_ conditions (**Figure 7B-C and Supplementary Figure S5**).

**Figure 7.**
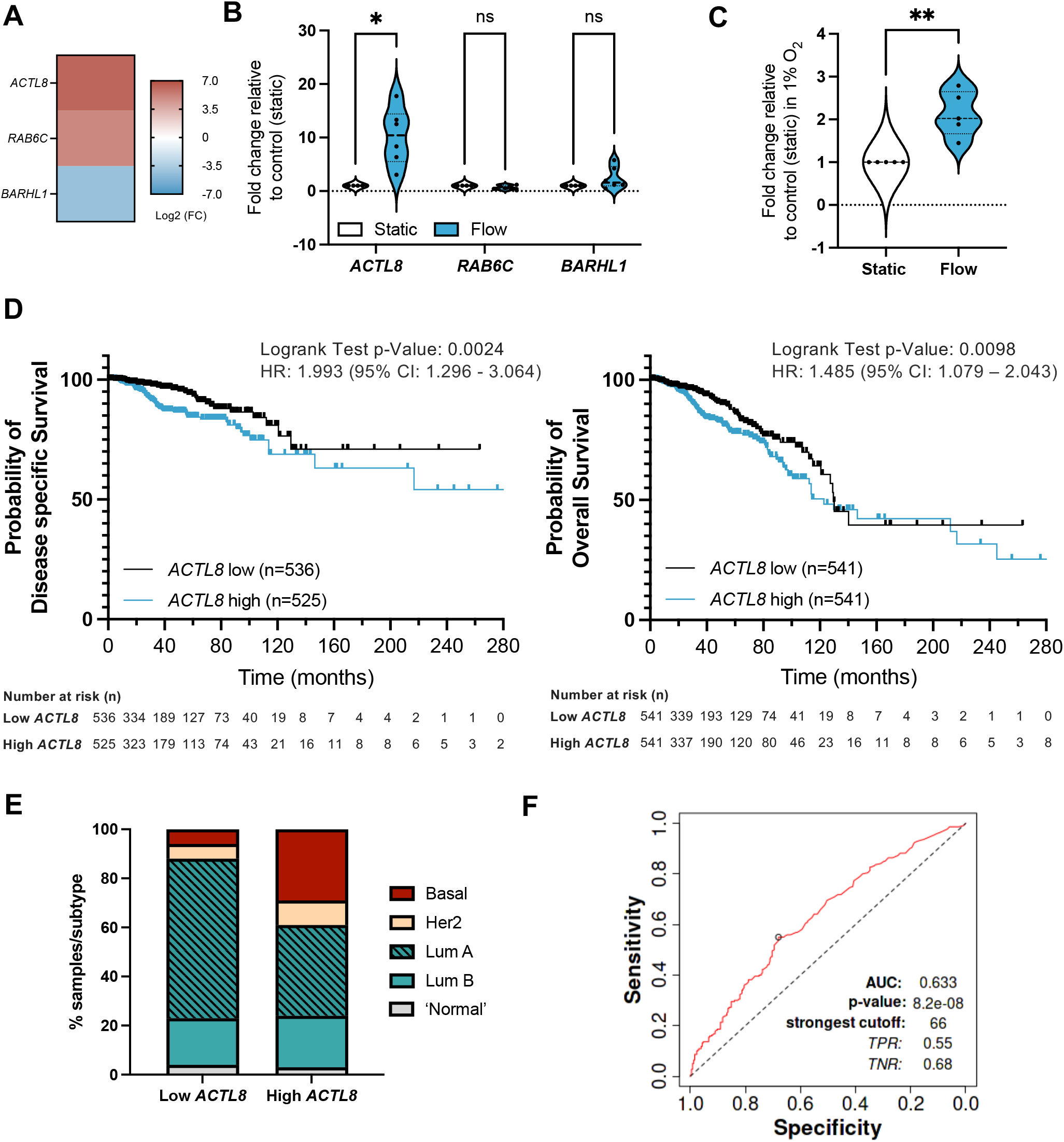
Identification of potential IFF-associated clinically relevant biomarkers – ACTL8 as a proof of principle analysis. (A-C) Panel of genes uniquely linked with either static or flow conditions were selected from the normoxic (20% O_2_) RNA-sequencing dataset, with Heatmap (A) representing log2 fold change for these genes. MCF7 spheroids were formed as before and exposed to static or flow conditions for 24 hours in the presence of Matrigel in normoxic conditions (20% O_2_), and gene expression for the gene panel validated through qPCR as described before, with plots (B) representing n=6 (24 h) independent experiments. Expression of ACTL8 was also evaluated in static or flow conditions for 24 h in the presence of Matrigel in hypoxic conditions (1% O_2_) (C). (D-E) Prognostic value for ACTL8 expression in breast invasive carcinoma patient samples was determined using the TCGA pan cancer Atlas RNA-seq dataset (n=1084) (Patient demographics in Supplementary Table 3). Disease-free Survival (n=1061) and Overall Survival (n=1082) plots (D) compare high and low ACTL8 expression, determined by median ACTL8 expression. This dataset was also analysed for proportion of breast cancer subtypes (E). (F) AUC (Area Under The Curve) ROC (Receiver Operating Characteristics) curve analysis was performed to evaluate the predictive value of ACTL8 gene expression vs responsiveness to any kind of chemotherapeutic treatment to determine predictive biomarker potential in breast cancer patients, n= 426 (non-responders = 197; responders = 229) as reported by Fekete and Gyorffy (2019) ^40^. For Survival plot analysis p value, a Logrank test was used; for ROC p value determination, a Mann-Whitney test was used. Significance is considered if p<0.05.

We performed *in silico* analyses of the prognostic value of ACTL8 in breast cancer patient samples (**Figure 7;** patient demographics in **Supplementary Table S3**). Survival analyses show that high expression of ACTL8 is a poor prognostic indicator of Disease-Specific Survival (time-period usually begins at the time of diagnosis or at the start of treatment and ends at the time of death), Overall Survival (overall patient survival status), Relapse-Free survival (time from treatment of disease to any event, irrespective of cause, except for any second primary cancers), and Distant Metastasis-Free Survival (Metastasis free status) (**Figure 7D and Supplementary Figures S6-S7**). We also observed that patient samples with high ACTL8 expression represent more aggressive breast cancer subtypes, including basal breast cancer, when compared to low-expressing patient samples (**Figure 7D**). ACTL8 had previously been identified as a potentially targetable cancer antigen for aggressive triple negative breast cancer (TNBC) ^59^, and linked with TNBC progression ^60^.

However, there is limited information on how its expression is regulated. *In silico* analysis of ACTL8 expression in breast cancer patient samples shows a positive correlation with hypoxia metagene signatures (**Supplementary Figure S8**), indicating ACTL8 as potentially regulated by hypoxia; whether this correlation is underpinned by a causal mechanistic relationship to hypoxia will be evaluated in further studies. TF enrichment analysis using Enrichr identified c-Myc as a potential transcriptional regulator of ACTL8 (p=0.02845, ChEA 2022 database). Expression of c-Myc has been shown to increase in response to shear stress, potentially downstream of YAP/TAZ via mechanosensing ^61-63^. It is therefore plausible to hypothesise that *ACTL8* could be a true c-Myc target gene in response to shear stress and IFF, but further work needs to be undertaken to validate this, alongside further unbiased analyses of TF binding to the ACTL8 promoter.

Interestingly, further overall survival analyses in other tumour types revealed that high expression of ACTL8 also has poor prognostic value, including for kidney, liver, head and neck, and endometrial cancer (**Supplementary Figure S9**), which again agrees with previously published work on head and neck cancer ^64, 65^.

Finally, to evaluate the biomarker value of ACTL8 in breast cancer we performed an ROC (Receiver Operating Characteristic) AUC (Area Under the Curve) analysis for a 5-year Relapse-Free Survival (RFS) for breast cancer patients who had received any type of chemotherapy and that were separated into responders (no residual histological tumour evidence after chemotherapy) or non-responders (patients with residual tumour tissue) (**Figure 7E**). For ACTL8, an AUC value of 0.63 indicates it has some predictive capacity for response to chemotherapy ^40^.

Therefore, these data indicate that ACTL8 could be a potential biomarker linked with interstitial flow for breast cancer and, potentially, other tumour types as well.

Regarding the role of ACTL8 in this context, previous studies have linked ACTL8 with PI3K/AKT signalling, which is known to contribute to proliferation, angiogenesis, and invasion ^60, 64^. However, prior to our study, no link with hypoxia or IFF has been identified. We hypothesise that potentially the biophysical TME mechanical cues linked with leaky vasculature, IFF, and hypoxia, drive the expression and likely role of ACTL8, but this remains to be evaluated.

ACTL8 has been described as an actin binding protein and a putative interactor with the dynactin complex via STRING analysis (**Supplementary Figure 10**), so it is therefore plausible that altered *ACTL8* expression could contribute to tumour progression via promotion of genomic instability, as reported for the dynactin complex ^66, 67^. To support this, we have performed additional analyses using the breast invasive carcinoma TCGA PanCancer dataset and have identified that high *ACTL8* expression is linked with increased genomic instability markers, such as fraction genome altered, *MYC* amplification, *TP53* mutations, and *PTEN* deletions (**Supplementary Figure 11**), but further experimental work needs to be performed to support these findings.

ACTL8 ultimately represents a case study for the use of genes whose expression is linked with IFF conditions as biomarkers. In the future, and analogous to hypoxia metagenes, more specific IFF gene expression signatures could be used as prognostic and predictive biomarkers.

## Conclusions

Although other studies have analysed transcriptional changes in high fluid flow rate/shear stress conditions in vasculature, such as those experienced by endothelial cells and circulating tumour cells ^8, 14^, our study using a spheroid on-chip microfluidic system is the first to systematically explore transcriptional changes linked with tumour IFF-like conditions, particularly in low oxygen conditions relevant to those observed in the hypoxic tumour microenvironment. Other studies have used microfluidic systems to model flow and oxygen levels and their impact on cell biology, particularly cell morphology and metastatic potential and drug responses ^68-72^. These primarily use media with varying oxygen conditions being flowed directly through the devices or oxygen depletion strategies directly on chip, rather than placing the whole system in a specialised hypoxia chamber, as in our study. Importantly, these studies focused on established signalling pathways, rather than using unbiased transcriptome-wide approaches and comprehensively compared these in normoxia vs hypoxic conditions to explore the underlying gene expression changes leading to biological changes underpinning these, as in our study.

As well as analysing broad gene expression and pathway enrichment changes, we have analysed ACTL8 as a candidate biomarker linked with IFF-like conditions and showed high expression of ACTL8 to be associated with poor prognosis in breast cancer. Identifying IFF-linked biomarkers would provide an alternative to traditional invasive and imaging-based methods to measure IFF, which have several limitations ^57^. Regions of hypoxia are linked with leaky vasculature and the occurrence of IFF conditions in tumours. However, the biological implications of these two biophysical TME factors remains poorly investigated, a gap which our model is addressing. It also allows us to evaluate the role of reperfusion and intermittent hypoxia and flow *in vitro* in controlled conditions, which has implications for response to therapy, as well as patient outcomes ^73^.

## Author Contributions

Conceptualization: EP, NP, IMP. Data curation: EP, MC, MR, IMP. Formal Analysis: EP, MR, MC, PRM, ST, IMP. Funding acquisition: AC, NP, IMP. Investigation: EP, MC, ST, IMP. Methodology: EP, MC, ST, AI. Resources: EP, MR, MC, ST, AI. Software: MR, MC. Supervision: AC, NP, IMP. Validation EP, PRM, ST, AI, NP, IMP. Visualization EP, MR, MC, AI, IMP. Writing – original draft: EP, IMP. Writing – review & editing: all authors

## Conflicts of interest

There are no conflicts to declare.

## Acknowledgements

EP was supported by a University of Hull PhD studentship. IMP, ST, MR, and AC are supported by Cancer Research UK RadNet Manchester [C1994/A28701]. PRM, AI, and NP are supported by Stockholm University. This project was further funded by a Biochemical Society Eric Reid award to IMP and University of Manchester & Stockholm University Joint Research Funding to IMP, NP, and AI.

## Notes

### Competing Interest Statement

The authors have declared no competing interest.

