## Supplementary Materials for "Investigating the impact of the interstitial fluid flow and hypoxia interface on cancer biology using a spheroid-on-chip perfusion system"

### Supplementary material

#### Supplementary Table S1 - List of primers used for qPCR in this study

| Target gene symbol | Catalogue reference or sequences 5' – 3' | Supplier |
|---|---|---|
| ***ABCA8*** | QT00011095 | Qiagen |
| ***ACTL8*** | QT00199934 | Qiagen |
| ***ASF1B*** | QT00007224 | Qiagen |
| ***B2M*** | QT00088935 | Qiagen |
| ***BARHL1*** | QT00214599 | Qiagen |
| ***BLM*** | QT00027671 | Qiagen |
| ***BNIP3*** | QT00024178 | Qiagen |
| ***BRCA1*** | QT00039305 | Qiagen |
| ***BRCA2*** | QT00008449 | Qiagen |
| ***CA9*** | F: GGAAGGCTCAGAGACTCA<br>R: CTTAGCACTCAGCATCAC | Sigma-Aldrich |
| ***CDH1*** | QT00019418 | Qiagen |
| ***E2F1*** | QT00016163 | Qiagen |
| ***EXO1*** | QT00080717 | Qiagen |
| ***FASN*** | QT00014588 | Qiagen |
| ***FEN1*** | QT00064722 | Qiagen |
| ***HK2*** | QT00013209 | Qiagen |
| ***LOX*** | F: GTTCCAAGCTGGCTACTC<br>R: GGGTTGTCGTCAGAGTAC | Sigma-Aldrich |
| ***LPIN1*** | QT00082873 | |
| ***MCM6*** | QT00059570 | Qiagen |
| ***MMP1*** | QT00014581 | Qiagen |
| ***MMP14*** | F: CCCCGAAGCCTGGCTACA<br>R: GCATCAGCTTTGCCTGTTACT | Sigma-Aldrich |
| ***PPARG*** | QT00029841 | Qiagen |
| ***PTGS2*** | QT00040586 | Qiagen |
| ***RAB6C*** | QT00219366 | Qiagen |
| ***RAD51*** | QT00031493 | Qiagen |
| ***RRM1*** | QT00066717 | Qiagen |
| ***RRM2*** | QT00039480 | Qiagen |
| ***SCD*** | QT00052381 | Qiagen |
| ***SNA1L*** | F: GACCACTATGCCGCGCTCTT<br>R: TCGCTGTAGTTAGGCTTCCGATT | Sigma-Aldrich |
| ***VEGFA*** | F: CTACCTCCACCATGCCAAGT<br>R: CTCGATTGGATGGCAGTAGC | Sigma-Aldrich |
| ***VIM*** | QT00095795 | Qiagen |
| ***WEE1*** | QT00038199 | Qiagen |
| ***WSB1*** | QT00064127 | Qiagen |

#### Supplementary Table S2 – Antibodies used in this study

| Target | Manufacturer | Reference or Catalogue No. | Dilution | Origin | Expected band size (kDa) |
|---|---|---|---|---|---|
| **β-Actin** | Santa Cruz | sc-47778 | 1:2000 | M | 42 |
| **E2F1** | Cell Signaling | | 1:1000 | | 70 |
| **E-cadherin** | Cell Signaling | 3195 | 1:1000 | R | 135 |
| **HIF-1α** | Becton Dickinson (BD) Biosciences | 610958 | 1:500 | M | 120 |
| **HK2** | Cell Signaling | 2867 | 1:1000 | R | 102 |
| **p53 (DO-1)** | Santa Cruz | sc-126 | 1:2000 | M | 53 |

M = mouse; R = rabbit

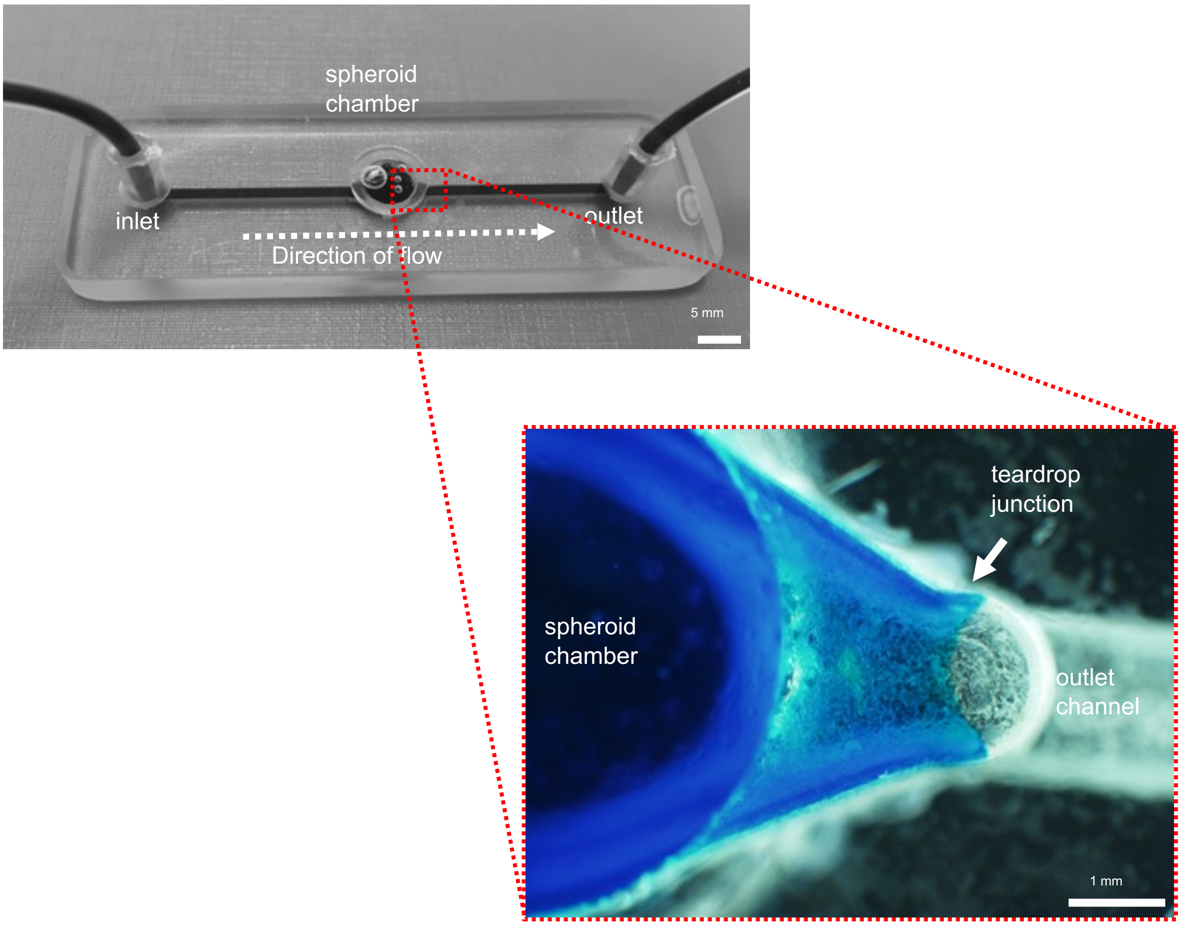


**Supplementary Figure S1 – Detail of teardrop junction in spheroid-on-chip perfusion device**
Photograph and microscope image detail of the spheroid-on-chip microfluidic perfusion device, highlighting the teardrop junction. Brilliant blue dye flowing through the chip demonstrates the location of the channel and shape of junction.

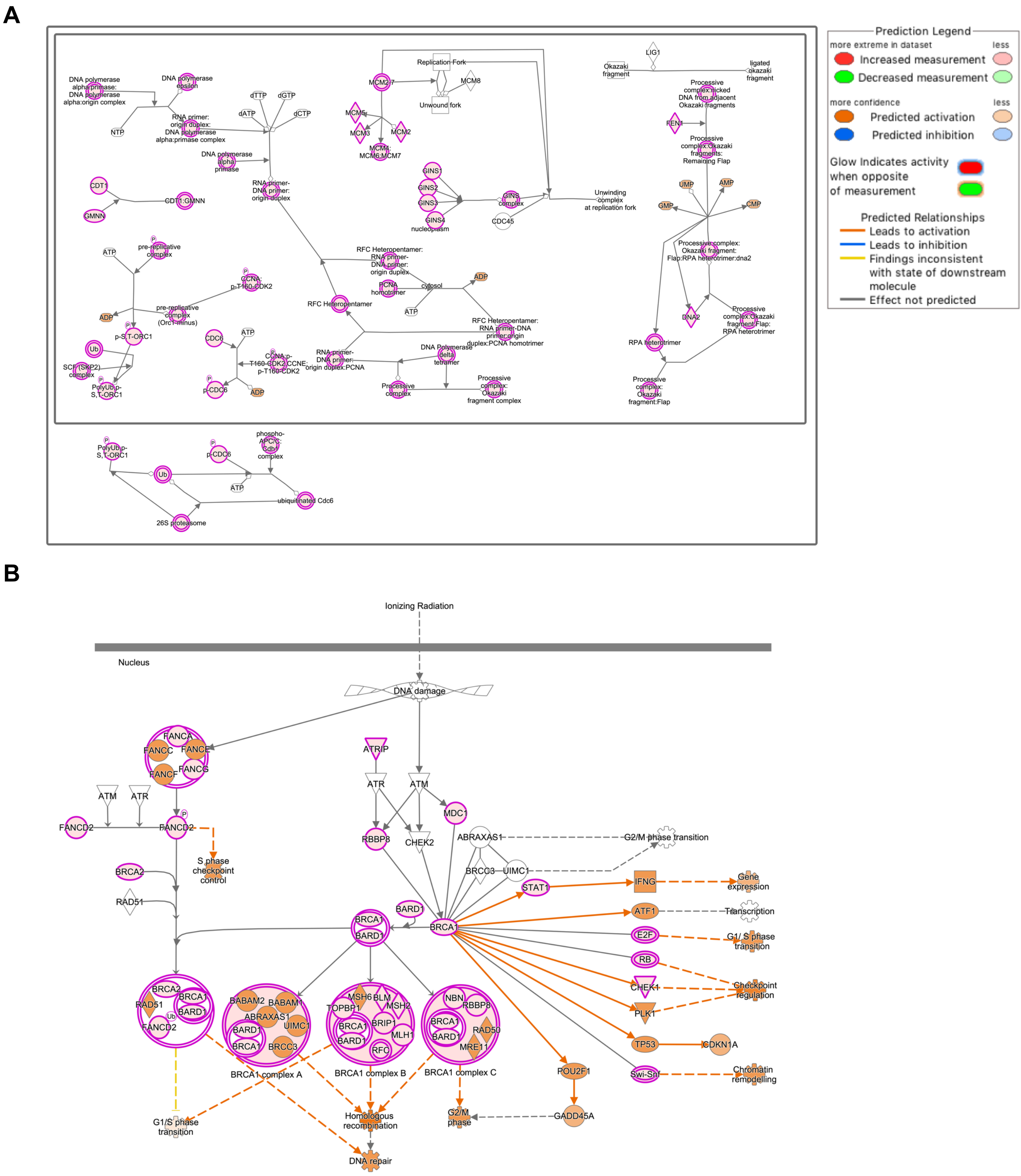


**Supplementary Figure S2 - IFF-like perfusion/flow led to upregulation of DNA replication, cell cycle, and DNA repair pathways**

Diagrams represent canonical pathway diagrams for DNA replication (A) and double strand break repair (B) generated using the Ingenuity Pathway Analysis platform.

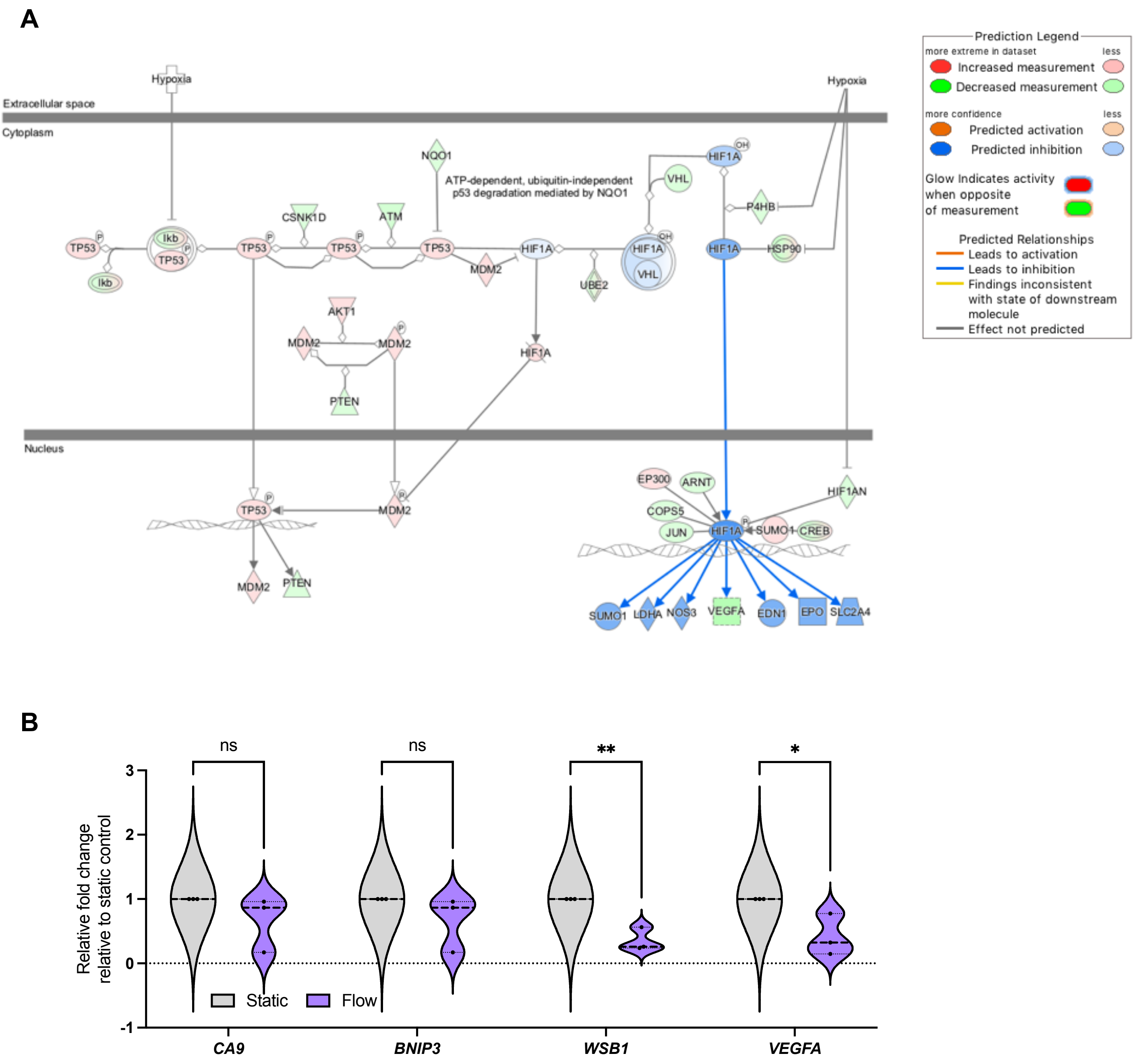


**Supplementary Figure S3 - Hypoxia signalling is downregulated in the better perfused spheroid-on-chip model**

(A) Canonical pathway diagram for low oxygen/hypoxia signalling generated using the Ingenuity Pathway Analysis platform.

(B) A panel of hypoxia-regulated genes was validated through qPCR as described before for 72 hours post flow, representing n=3 independent experiments. Unpaired student's t-test was performed to test for statistical significance between samples. ns = nonsignificant; * $p<0.05$; ** $p<0.01$

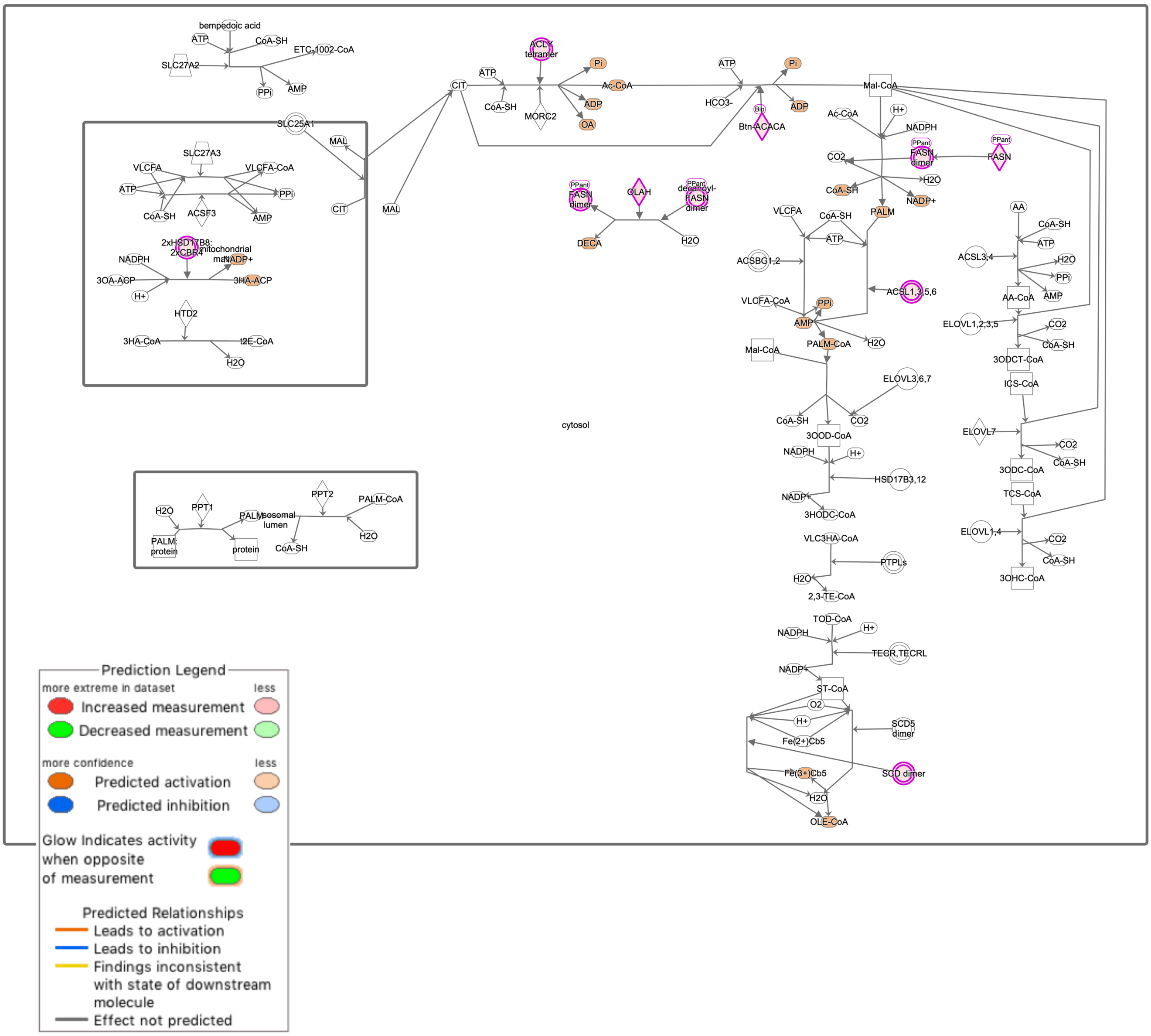


**Supplementary Figure S4 - Gene expression changes mediated by IFF-like perfusion/flow differ depending on environmental oxygen tensions**

Diagram represents canonical pathway diagrams for Phospholipid metabolism signalling pathways were enriched in upregulated DEGS after flow generated using the Ingenuity Pathway Analysis platform.

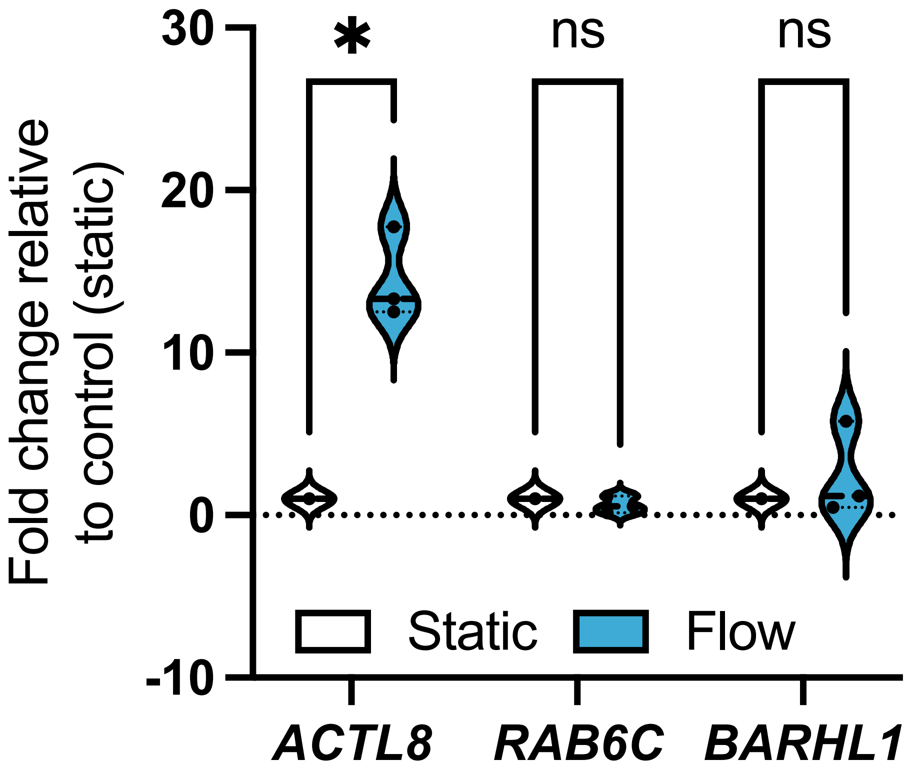


**Supplementary Figure S5 – Identification of potential IFF-associated clinically relevant biomarkers – validation of candidate panel at 72 hours**

A panel of genes uniquely linked with either static or flow conditions were selected from the normoxic (20% $O_2$) RNA-sequencing dataset (as determined in Figure 2). MCF7 spheroids were formed as before and exposed to static or flow conditions for 72 hours in the presence of Matrigel in normoxic conditions (20% $O_2$), and gene expression for the gene panel validated through qPCR as described before, with plots (B) representing n=3 independent experiments.

**Supplementary Table S3 - Patient demographics of breast invasive carcinoma TCGA pan cancer Atlas dataset**

| | | |
|---|---|---|
| **Age**<br>median (range) | 58 (26 - 90) | |
| **Sex**<br>n (%) | *Female* | 1072 (99) |
| | *Male* | 12 (1) |
| **Disease Stage ***<br>n (%) | I | 189 (17) |
| | II | 613 (57) |
| | III | 244 (22) |
| | IV | 19 (2) |
| | Unknown | 19 (2) |
| **Molecular Subtype**<br>n (%) | Basal | 171 (16) |
| | Her2 | 78 (7) |
| | Luminal A | 499 (46) |
| | Luminal B | 197 (18) |
| | 'normal' | 36 (3) |
| | Unknown | 103 (10) |

*Neoplasm Disease Stage American Joint Committee on Cancer Code

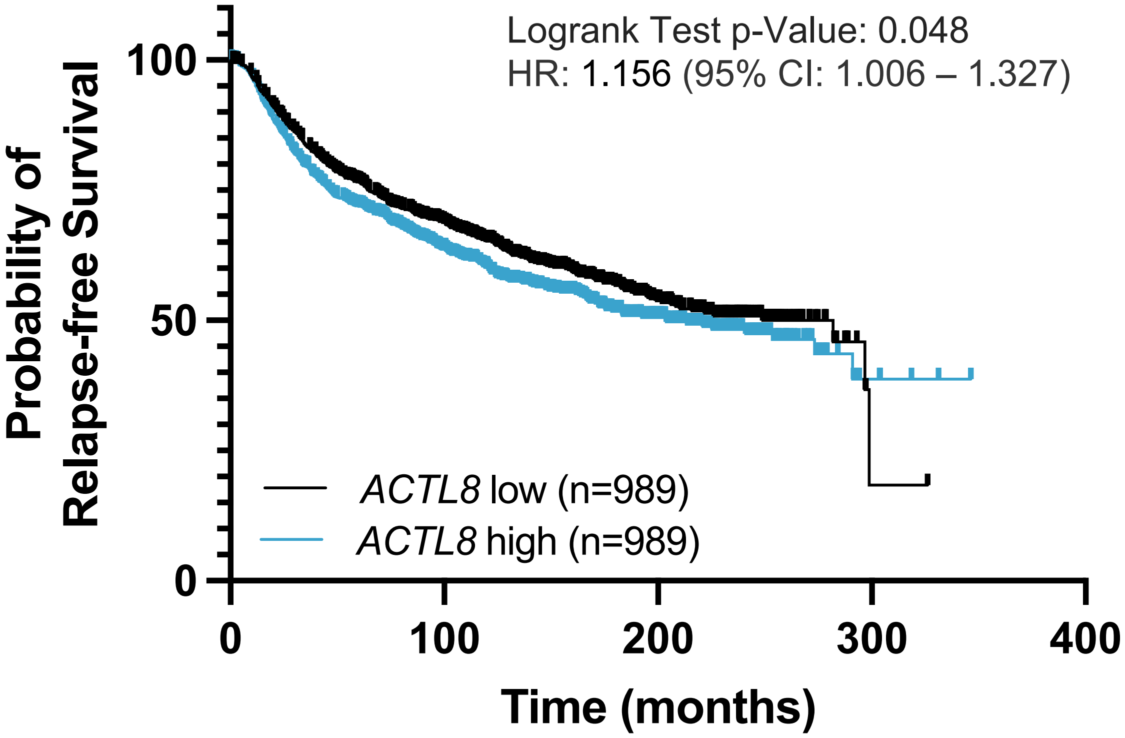


**Supplementary Figure S6 – Prognostic value for *ACTL8* expression in breast cancer patient datasets (Metabric Dataset)**

Breast invasive carcinoma patient samples was determined using the Metabric RNA-seq dataset (n=1978). Relapse-free Survival plot compare high and low *ACTL8* expression, determined by median *ACTL8* expression.

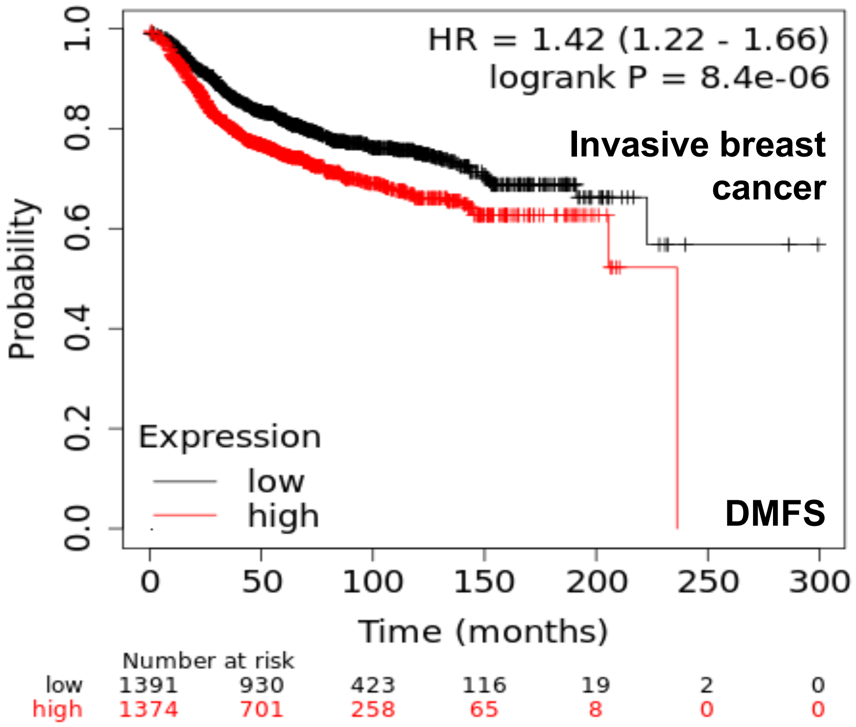


**Supplementary Figure S7 – Distant Metastasis-Free Survival prognostic value for *ACTL8* expression in breast cancer patient datasets**
KMplotter was used to analyse *ACTL8* expression levels in gene chip array-based patient data and correlate it to distant metastases-free survival (DMFS) *n* =2765. Patients were split by *ACTL8* median expression, and no other patient selection thresholds were applied.

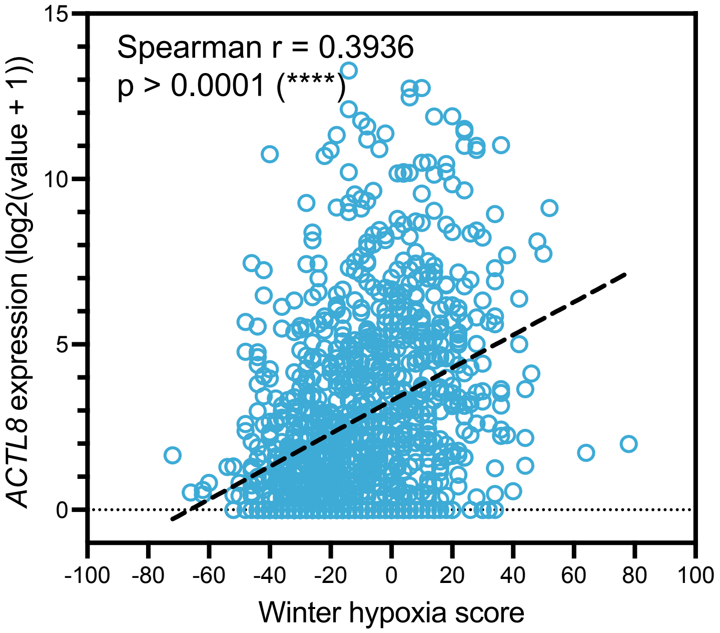


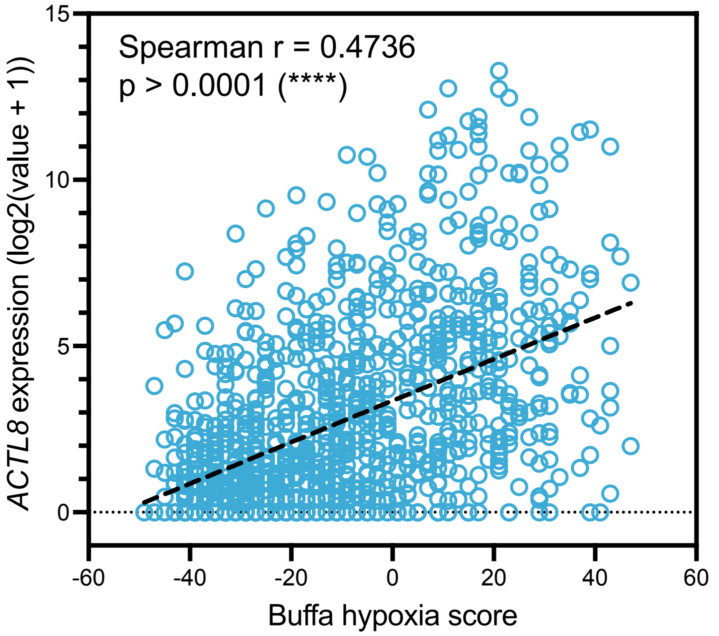


**Supplementary Figure S8 – *ACTL8* expression in relation to hypoxia scores**
*ACTL8* expression in patients ($n$ = 994) from the TCGA PanCancer Atlas Breast Invasive Carcinoma study is plotted on the y-axis against the Winter and Buffa hypoxia score. The hypoxia score is representative of a hypoxia metagene signature. The expression data was log2 transformed.

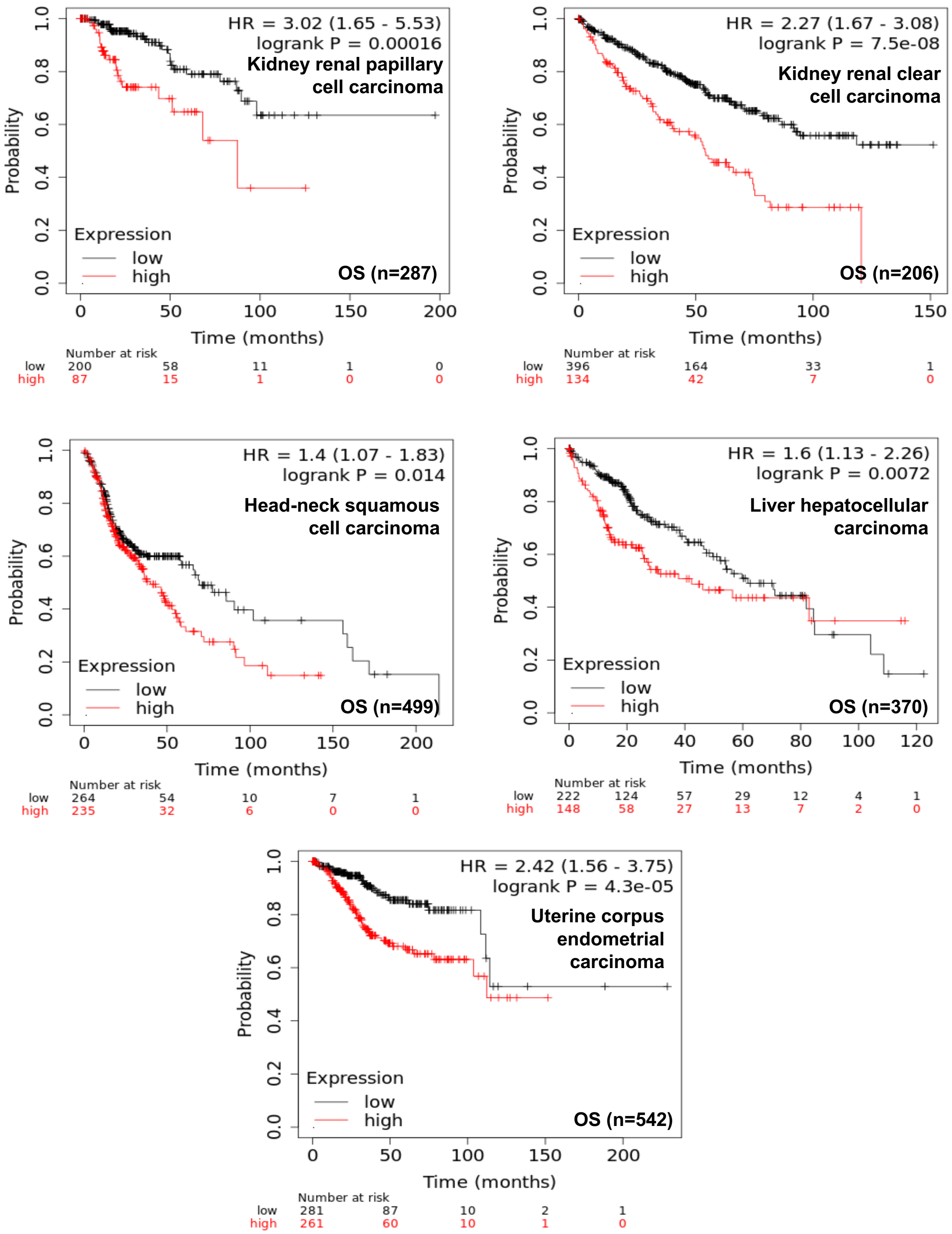


**Supplementary Figure S9 – Overall Survival prognostic value for *ACTL8* expression in other tumour types (pan-cancer analysis)**

KMplotter was used to analyse *ACTL8* expression levels in gene chip array-based patient data and correlate it to Overall Survival. Patients were split by *ACTL8* median expression, and no other patient selection thresholds were applied.

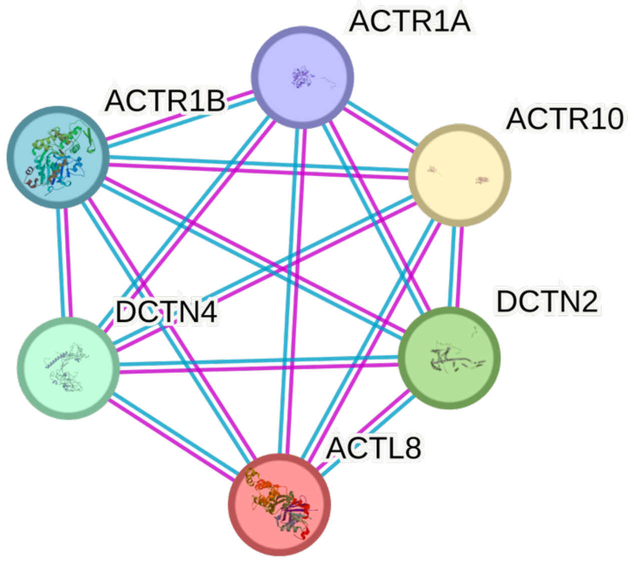


**Supplementary Figure S10 – STRING PPI (protein-protein interaction) analysis network for ACTL8**

The interactome was produced using the online STRING database (Szklarczyk et al., 2015). Colour of the connectors depends on the type of evidence for the interaction: blue: from curated databases; pink: experimentally determined.

Key: ACTL8 (Actin like 8); DCTN2 (Dynactin subunit 2); DCTN4 (Dynactin subunit 4); ACTR1A (Alpha-centractin); ACTR1B (Beta-centractin); ACTR10 (Actin related protein 10)

**A**

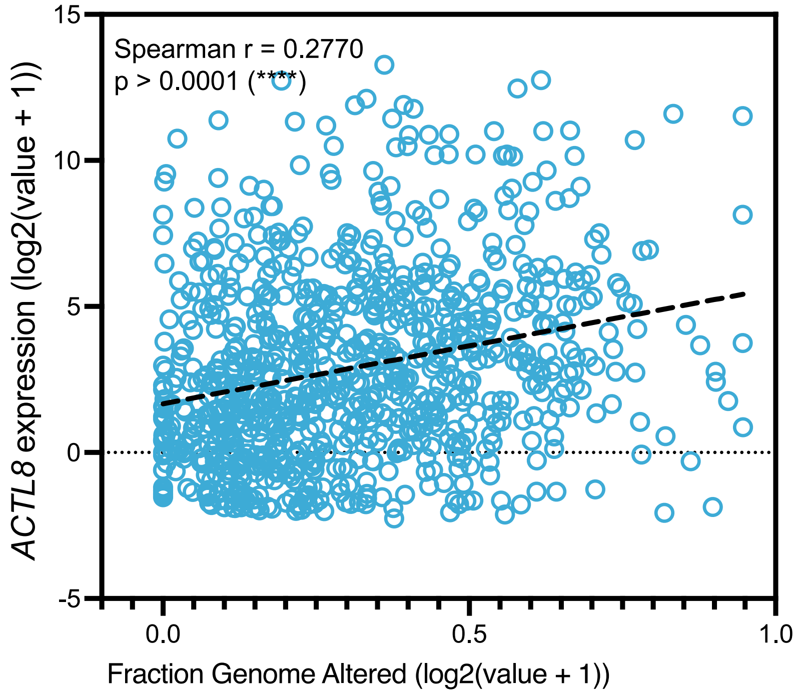


**B**

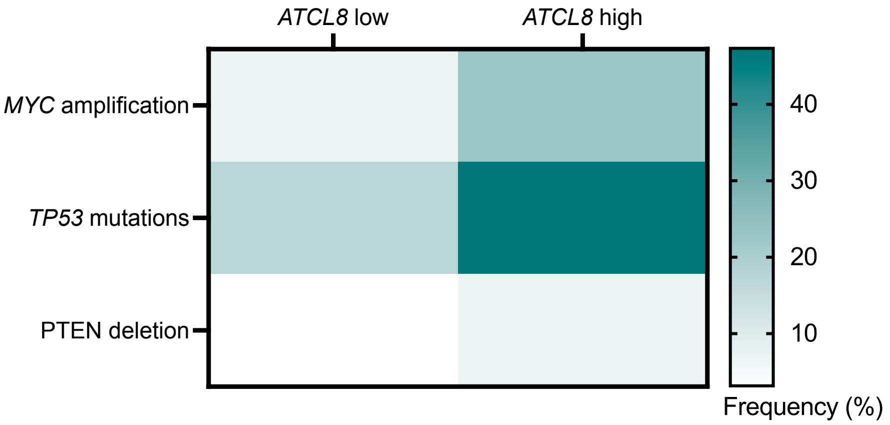


**Supplementary Figure S11 – Analyses of relationship between ACTL8 mRNA expression and genomic instability markers**

*ACTL8* expression in patients (*n* = 994) from the TCGA PanCancer Atlas Breast Invasive Carcinoma study were analysed for links with genomic instability markers.

A - Correlation analysis of *ACTL8* mRNA levels vs Genomic instability marker ‘Fraction Altered Genome’.

B - Frequency of genomic alterations linked with increased genomic instability (*MYC* amplification, *TP53* mutations, and *PTEN* deletions) comparing patient samples with low of high *ACTL8* mRNA expression.

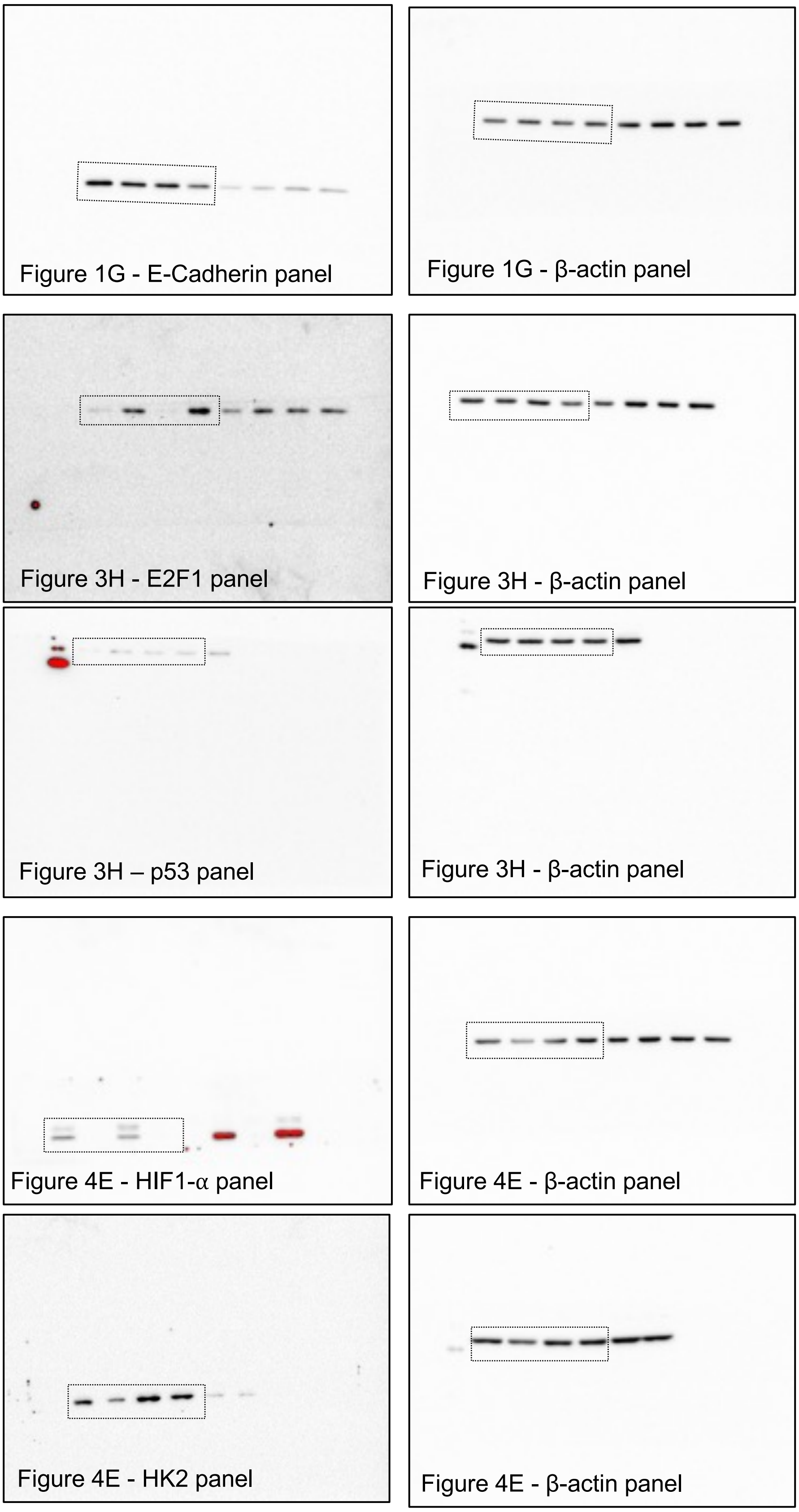


**Supplementary Figure S12 – Raw blots from immunoblotting panels in manuscript**
